# Automated Workflows using Quantitative Colour Pattern Analysis (QCPA): A Guide to Batch Processing and Downstream Data Analysis

**DOI:** 10.1101/2023.02.02.526788

**Authors:** Cedric P. van den Berg, Nicholas D. Condon, Cara Conradsen, Thomas E. White, Karen L. Cheney

## Abstract

Animal and plant colouration presents a striking dimension of phenotypic variation, the study of which has driven general advances in ecology, evolution, and animal behaviour. Quantitative Colour Pattern Analysis (QCPA) is a dynamic framework for analysing colour patterns through the eyes of non-human observers. However, its extensive array of user-defined image processing and analysis tools means image analysis is often time-consuming. This hinders the full use of analytical power provided by QCPA and its application to large datasets. Here, we offer a robust and comprehensive batch script, allowing users to automate many QCPA workflows. We further provide a set of useful R scripts for downstream data extraction and analysis. These scripts will empower users to exploit the full analytical power of QCPA and facilitate the development of customised semi-automated workflows. Such quantitatively scaled workflows are crucial for exploring colour pattern space and developing ever-richer frameworks for analysing organismal colouration accounting for the visual perception in animals other than humans. These advances will, in turn, facilitate testing hypotheses on the function and evolution of vision and signals at quantitative and qualitative scales, which are otherwise computationally unfeasible.

## 1. Introduction

Understanding the perception of visual information by non-human observers is crucial to studying the ecology and evolution of animal and plant colouration. The last two decades have seen the creation and widespread adaptation of tools and methods that allow researchers to simulate or approximate aspects of animal vision, such as colour contrast perception (e.g. Endler and Mielke, 2005; Gawryszewski, 2018; Kemp et al., 2015; Renoult et al., 2017; Vorobyev and Osorio, 1998) and spatial vision (Godfrey et al. 1987; Caves and Johnsen 2018). These advances coincide with the steady development of colour pattern analyses (e.g. Chan et al., 2018; Endler, 1991; Endler, 2012; Stoddard et al., 2014; Van Belleghem et al., 2018; van den Berg et al., 2020) and their integration into increasingly comprehensive collections of tools and functions across software platforms such as pavo (Maia et al. 2019) in R (R Core Team 2021) or the Multispectral Image Calibration and Analysis toolbox (MICA) (Troscianko and Stevens 2015) in ImageJ (Schneider et al. 2012).

Quantitative Colour Pattern Analysis (QCPA) (van den Berg et al. 2020b) is a recent and powerful addition to this landscape. It provides a dynamic analytical framework for analysing visual scenes through the eyes of ecologically relevant observers and is integrated into the MICA toolbox (Troscianko and Stevens 2015). It allows users to choose preferred tools and analyses uniquely suitable for analysing spatiochromatic information. Briefly, it achieves this by taking full-spectrum images as its input (thereby generating ‘.mspec’ files) before applying a suite of models which can consider the spectral, spatial, and temporal sensation of non-human viewers, to ultimately give image-based and numerical outputs that summarise the structure of visual scenes and stimuli (van den Berg et al. 2020b).

Preparing and processing calibrated images in the framework, however, remains tedious. This is due to the time needed to translate calibrated digital images into cone catch images, user-guided input for identifying regions of interest (ROIs), and the subsequent application of image processing and analyses. Saving and labelling multiple output files, as well as extracting data for subsequent statistical analysis, also remains a tiresome process. As a result, applying the QCPA at quantitative scales, such as analysing hundreds or thousands of images at multiple viewing distances, is nearly impossible in its original form.

An image currently needs hours of manual, repetitive work to obtain the full numerical output from the various types of analyses in the QCPA framework. This is especially true when considering multiple viewing distances and regions of interest (ROIs). Consequently, most studies using the framework do not consider more than a few dozen individual observations with a limited subset of available image statistics (e.g., Nokelainen et al., 2021; Rodríguez-Morales et al., 2021). This is problematic as quantifying ecologically relevant effects, such as colour pattern variability in natural populations or the analysis of behavioural data, requires sufficiently large sample sizes to provide adequate statistical power. This holds particularly true for a method producing hundreds of image statistics from a single observation, such as the QCPA, where the number of available image statistics easily outnumbers observations (van den Berg et al. 2022). Considering all possibly relevant image statistics and deducing a relevant set of parameters is often the desired approach instead of pre-emptively narrowing down the number of considered image statistics (van den Berg et al. 2022). However, such deductive approaches require adequate statistical solutions and remain an intriguing challenge in using complex colour pattern spaces (Stoddard and Osorio 2019; van den Berg et al. 2020b, 2022). The QCPA framework is, therefore, in urgent need of automation to facilitate its complete application to large datasets.

Here, we present a dynamic batch-processing extension to QCPA, allowing users to apply almost the entire QCPA framework or selected parts to large datasets via a flexible, GUI-guided input. We further provide a set of R scripts that efficiently extract numerical data from the consecutively stored output files for downstream analysis. We describe the functionality of both—the QCPA batch script and complementary R scripts—and provide detailed worked examples. While this script does not provide a complete solution for every possible application of the QCPA, we hope it will offer a viable solution to most. This batch script should also help researchers customise their automated pipelines, stimulating the open exchange of programming solutions via platforms such as open-access publications, open-access platforms such as GitHub or the dedicated user forum for the MICA toolbox (www.empiricalimaging.com).

## 2. QCPA batch script

### 2.1. Intended use

The QCPA batch script is intended to quantify spatiochromatic information of an animal against its visual background. As such, the script can also be applied to images without an animal or object of interest or an animal or object itself. The batch script is intended for users with moderate experience in the manual application of QCPA.

The script allows the user to choose the following desired analytical outputs individually: 1) Colour Adjacency Analysis (CAA), Visual Contrast Analysis (VCA), Boundary Strength Analysis (BSA) and particle analysis; 2) Local Edge Intensity Analysis (LEIA); 3. Colour Maps; and 4) GabRat. For a detailed discussion of these analyses, please see the original publications and their modifications for QCPA as listed in van den Berg et al. (2020).

Notably, the script requires cone mapping functions derived from calibrated cameras (several standard profiles included in the MICA toolbox) and currently does not permit the use of chart-based models. While covering many analyses available in QCPA and the MICA toolbox, the batch script extension does not provide a comprehensive library of automation. However, the script is designed to be easily modified by the user. Such modifications are explicitly invited from the community, and we encourage their open sharing on community platforms in the interest of mutual benefit among researchers in the field.

### 2.2. Data preparation

Image processing in QCPA is specific to a variety of user choices specifying the properties of the observer’s visual system, the light environment in which a picture was taken as well as the intended target light environment, observer viewing distance, and a variety of parameters and processing choices that vary depending on the desired analyses. The batch script uses three approaches to minimise the need to repeat the number of times such input is required:

1. Pre-defined folder structure
2. Auxiliary files
3. Batch processing graphical user interface (Batch GUI)

#### 2.2.1 Folder structure

The batch script requires input data to be organised in a specific way. This refers to the naming of files and the folder structure. For example, the script expects individual observations (i.e., animals or objects) to be grouped within an overarching folder (i.e., species or site) and file naming must be uniform. This allows the script to reliably detect individual observations while allowing the user to structure their data according to sites or taxa (see the provided datasets for specific examples).

#### 2.2.2 Auxiliary files

The batch script uses the MICA toolbox’s pre-existing approach to associate several files with each image to be analysed. In addition to the already existing need for a folder with the specified ROIs and the corresponding .mspec of each image, the batch script requires two additional text files specifying the rotation of each image and the cone mapping function (see the manual for details). These allow users to standardise the orientation of all images before analysis and permit multiple cone mapping functions within the same dataset (e.g., multiple lighting environments).

#### 2.2.3. Batch GUI

The batch script allows the user to specify which of the available analyses they want to conduct and what settings they want to use. The Batch GUI consists of a central interface and a suite of dedicated, tool-specific GUIs activated by user choices. These tool-specific GUIs provide helpful information and allow running all analyses with independent settings. Furthermore, the batch script remembers the most recent user input and will pre-fill the previous choices by the user, helping to test settings and maintain track of chosen settings in case of incomplete runs. See the manual and worked examples for detailed instructions on how to use them. Users familiar with QCPA and MICA toolbox will find many GUIs similar to existing ones. However, the batch script features several custom-built GUIs to enable the flexible use of various user-defined processing steps.

### 2.3. Workflow recommendations

The Batch GUI, manual and worked examples contain a suite of recommendations in addition to those in van den Berg et al. (2020) and the empiricalimaging.com website that users might find informative. We recommend that the user validate the numerical accuracy of the batch script output on a small dataset before analysing large datasets. This also allows the user to confirm that the folder structure and auxiliary files have been arranged correctly. As processing large datasets can take many hours, days, or even weeks, we recommend that the user consider running sub-sections of larger datasets on different instances, such as multiple computers, servers, or cloud-based computing facilities. This will ultimately save time and prevent data loss while helping to detect faulty files in large datasets faster.

Like all approaches to visual modelling, image analysis with QCPA requires many different input choices by the user. Keeping a record of these choices is crucial for two reasons. First, it allows the user to keep track of settings, facilitating record-keeping and collaboration. Second, it provides for the publication of repeatable research. The latter is often an issue in visual modelling studies, as modelling choices are often poorly documented (White et al. 2015). To this end, the QCPA batch script keeps various detailed log files in the data output. These can easily be added to the supplementary information of any publications or uploaded/shared as part of the data.

## 3. R script library

We provide a set of tailored R-scripts enabling the extraction and compilation of batch QCPA output data into .csv tables that can be used for downstream analysis. The efficient handling of QCPA batch output is crucial in using QCPA at larger scales, as various outputs are stored in files across a dedicated data set structure. The scripts and functions we provide here are formulated and commented on in a way that is aimed to facilitate modification by the user. These scripts and functions are intended to provide guidance for people with limited experience in managing large and complex datasets. However, these scripts and functions also aim to provide a starting point for more versed users to tailor their own data analysis pipelines. A detailed manual for using the R scripts can be found in the electronic Supplement or here: https://github.com/CaraConradsen/QCPA-r-script.

The R environment provided includes the following functionality:

1. Compiling CAA, VCA, BSA, and particle analysis data.
2. Compiling data from LEIA
3. Compiling data from GabRat
4. Compiling data from Particle Analysis and secondary data output
5. Merging, data cleaning and reshaping various subsets of data
6. Exploratory data visualisation and analysis
7. Exporting compiled files

## 4. Worked examples

We provide detailed worked examples with test data and corresponding toolbox files, such as cone mapping models. These aim to highlight the functionality of the batch script and allow users to familiarise themselves with examples related to their needs. Instructions for the worked examples can be found in the electronic Supplement or here: https://github.com/cedricvandenberg/QCPA-batch-script.

### 4.1 Trichromatic example without UV (sea slugs seen by a triggerfish, *Rhinecanthus aculeatus*)

Here, we showcase the use of the batch script by applying all modules in the script to investigate how a triggerfish (*Rhinecanthus aculeatus*) perceives two species of cryptic sea slugs (*Aphelodoris varia, Aplysia sp*.) from the east coast of Australia (Nelson Bay, New South Wales) photographed underwater (Fig. 1).

**Figure 1.**
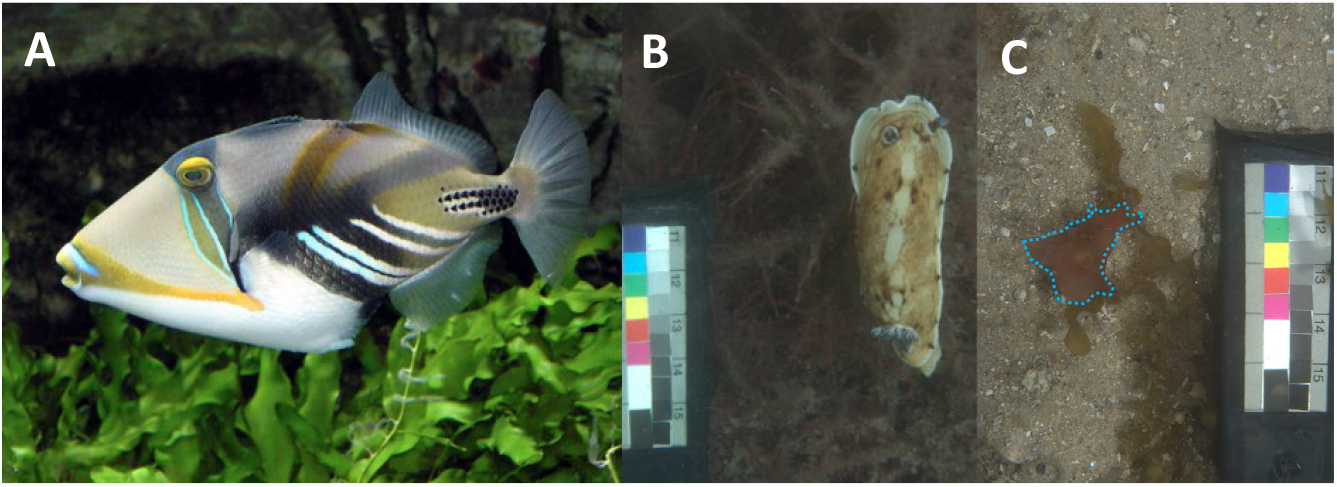
A) *Rhinecanthus aculeatus* (A. Pingstone, public domain). Calibrated images from the test dataset of *Aphelodoris varia* (B) and *Aplysia sp*. highlighted with blue dashed line (C) by CvdB.

### 4.2 Tetrachromatic example with UV (spiders seen by a bluetit, *Parus major*)

This example uses all batch script modules to analyse cryptic spiders (*Tamopsis brisbanensis*) found on eucalypt trees in Sydney, NSW (Australia) (Fig. 2).

**Figure 2.**
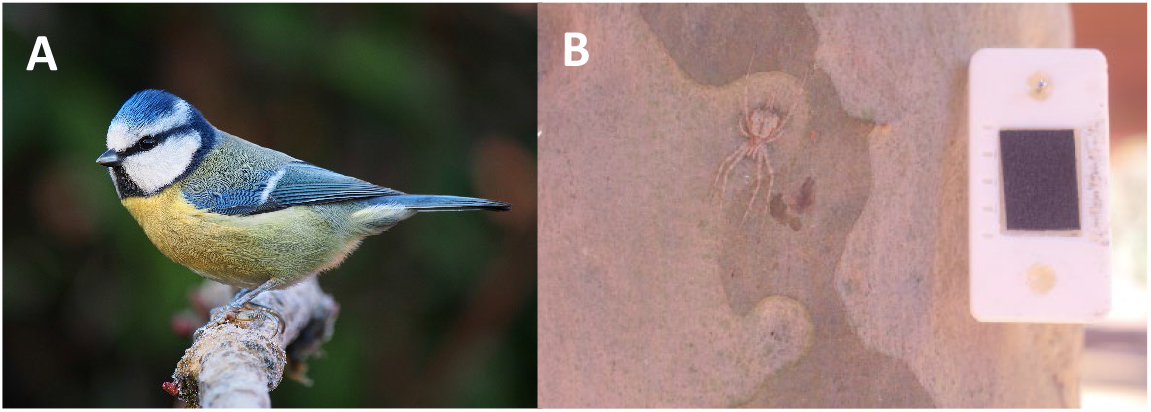
A) *Parus major* (© Francis C. Franklin / CC-BY-SA-3.0); B) Calibrated image of *Tamopsis brisbanensis*, taken by A. Aceves (with permission).

## Discussion

We provide a user-friendly, open-source batch processing extension to QCPA, allowing for the semi-automated analysis of large datasets within the QCPA framework. This greatly facilitates the full-scale use of the analytical power of the framework as intended by its original design (van den Berg et al. 2020b). The open-source code of the script is presented and structured with the deliberate purpose of enabling users to create customised versions of the batch script, which might be more suitable to their intended purposes. The batch script comes with a detailed manual guiding users in preparing and analysing the image data. We further provide three worked examples and corresponding test data, covering many contexts where users will likely use the script. Lastly, the user is provided with a set of R-scripts aiding in the downstream analysis of the data generated by the batch script. These scripts are presented and structured in a way that facilitates users to customise them to their needs.

The script enables the semi-automated analysis of large image datasets, computing hundreds of colour pattern descriptors at multiple viewing distances for dozens, hundreds, or even thousands of observations. This has previously been impossible due to the time-consuming manual processing of images in QCPA. As QCPA and other colour pattern analyses using visual modelling approaches (e.g. Maia et al., 2019; Troscianko and Stevens, 2015) merely relate to approximations of early-stage visual processing (e.g. Endler and Mielke, 2005; Marr, 2010; van den Berg et al., 2020a; van den Berg et al., 2020b; Vorobyev and Osorio, 1998), considering the context and task-dependent relationships with and between a broad range of descriptors is critical (see van den Berg et al., 2022 for discussion). Thus, the presented batch script provides a crucial step towards such analytical approaches.

Unlike other image analysis tools available to visual ecologists, such as pavo (Maia et al. 2019), MICA is an ImageJ (Schneider et al. 2012) plugin written in Java (Arnold et al. 2005). It is intentionally tailored towards accessibility via graphical user interfaces (GUIs) rather than command line prompts. Unlike R software (R Core Team 2021) or Python (van Rossum 1995), frequently taught in undergraduate and HDR Biology degrees worldwide, Java is not a programming language that biologists are readily familiar with. Therefore, writing scripts and macros for a complex network of GUI-guided tools in an unfamiliar language is a hurdle for most users of the QCPA framework seeking to batch-process large volumes of image data. This creates the potential for unintentional gatekeeping by researchers with access to programmers familiar with writing scripts in Java and an adequate understanding of the framework’s mechanisms. The result is a hidden custom of informally traded scripts, which undermines fast and equal access to methods in the field.

QCPA provides a conceptual approach towards combining visual modelling and colour pattern analyses. It does not claim to have a ‘perfect’ way of achieving its purpose. Rather, it provides an idea of how to combine different elements of vision modelling and colour pattern analysis, lending from and contributing to existing methods. Therefore, automating workflows and advancing customisability represent essential stepping stones towards the continued development and refinement of the framework and its contribution towards alternative methods. By providing an inviting, well-annotated, intuitive, open-access and open-source environment, we hope to facilitate the application of and, importantly, eventual modifications and additions to the framework and its combined use with other existing methodologies. Cumulatively, this round-table philosophy will not only advance the development and testing of methods in the field but, crucially, will facilitate the increased critical understanding of, and familiarity with, existing methodology among researchers in visual ecology. On a more tangible note, this script can easily be adjusted to enable the analysis of different data structures or include processing steps and analyses available in QCPA or the MICA toolbox currently not considered in the script. The R scripts and functions provided here provide useful guidance for users with variable experience in data wrangling. However, more extensive forms of such customised data extraction functions will provide an important step towards the dynamic use of large-scale QCPA output across multiple colour pattern analysis frameworks in the R environment, such as pavo.

## 6. Acknowledgements

We thank Alfonso Aceves for providing the test data for the UV examples.

## 7. Declarations

### 7.1 Funding

This work was supported by the Australian Research Council (FT190199313 to KLC), Hermon Slade Foundation grant to TW (HSF20082), and a UQ Research Stimulus (Allocation Two) Fellowship as well as a Swiss National Foundation Postdoc.Mobility Research Fellowship (P500PB_211070) awarded to CvdB. NC was supported as a CZI Imaging Scientist by grant number 2020-225648 from the Chan Zuckerberg Initiative DAF, an advised fund of the Silicon Valley Community Foundation.

### 7.2 Conflicts of interest

The authors declare no conflict of interest.

### 7.3. Ethics approval

Not applicable

### 7.4. Consent to participate

Not applicable

### 7.5. Consent for publication

All authors consent to the publication of the manuscript and associated documents.

### 7.6 Availability of data and material

All data and supplementary materials are available at the respective repositories listed below.

### 7.7 Code availability

All code is available at the respective repositories listed below.

## 8. Author’s contributions

CvdB – Overall concept (lead), testing, coding, original draft, manuscript writing & review, Funding, NC –coding batch script (lead), Manuscript review, CC – R scripts (lead), Manuscript review, TEW – R scripts, Manuscript review, Funding, KLC – Original concept, Funding, Manuscript review.

## 9. Supplementary data

Batch script ImageJ plugin, Batch script manual, worked examples guide: https://github.com/cedricvandenberg/QCPA-batch-script

R scripts and functions:

https://github.com/CaraConradsen/QCPA-r-script

Test data for the worked examples:

https://doi.org/10.48610/3cdcc1f

## Notes

https://github.com/cedricvandenberg/QCPA-batch-script

https://github.com/CaraConradsen/QCPA-r-script

https://doi.org/10.48610/3cdcc1f

### Competing Interest Statement

The authors have declared no competing interest.

### Summary of Updates

Various typos and final edits prior to submission with Evolutionary Ecology

