## Supplementary material for "Automated Workflows using Quantitative Colour Pattern Analysis (QCPA): A Guide to Batch Processing and Downstream Data Analysis": Manual (from another version of this preprint: version 2, 2023-11-10)

### 1. General information

This script is targeted toward quantifying an animal and its visual background as well as an entire visual scene. The presented workflow is designed for RAW files processed with calibrated cameras, using visual models.

Note: It is not possible to use .mspec files generated using calibrated jpgs with this script.

This script is an advanced application of the QCPA framework, and we recommend the user get familiar with the manual workflow first. We further recommend testing the script and the accuracy of numerical output on small subsets of larger data sets and their manually obtained output.

This script has been developed using Windows and Linux-based systems but should work well on Macs.

If using this script, cite the pre-print (available here) in addition to the original QCPA & MICA papers (available here).

### 2. Installing the batch script

Extract the ‘Batch QCPA.zip’ folder and paste it into the ‘micaToolbox’ folder.

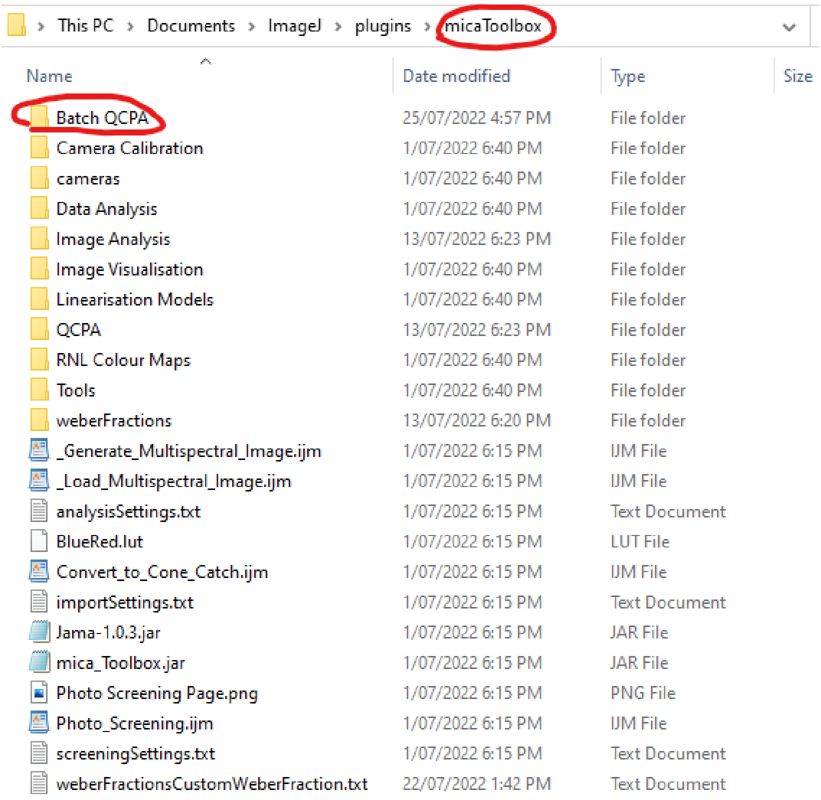

#### 3. Required folder structure

The script requires a specific folder structure. Specifically, individual observations (e.g. an animal on its background, a scene, etc.) need to be nested in an 'individual' folder within a 'site' or 'species' folder. The script is then designed to work on a single folder containing this simple structure. See the worked examples data structure for specific examples.

#### 4. Auxiliary files

In addition to the .mspec and the original RAW image file(s), the script requires a specific set of auxiliary files in each folder. Namely:

1) a .zip file containing the regions of interest (ROIs) (mandatory)
2) a .txt file with the desired image rotation (optional)
3) a .txt file specifying the visual model to be used (mandatory)

It is easiest to obtain these files in the following order for each image:

#### 4.1 Specifying the image rotation

Colour pattern analyses tend to be very sensitive to image orientation. We therefore highly recommend standardising images to a uniform orientation prior to analysis. To do so, after selecting 'Generate multispectral image' and selecting your appropriate settings, go to 'image'-'Transform'-> 'rotate…' Select 'Preview' and 'Zoom to fit'. Align the animal head up using the gridlines displayed. Make sure nothing gets clipped. Save the rotation angle as a .txt called 'rotation' with the image.

#### 4.2 Specifying the visual model

The batch script allows for multiple visual models to be used with a given dataset. For example, if the dataset contains images that were shot during sunset and images that were shot during broad daylight and yet other images which were shot in forest shade. Provided the user has measured these light environments a set of lighting-specific models can be trained using the 'Generate cone catch model' function in the 'Camera calibration' menu. The user can then place a .txt file called 'cone model' with each picture specifying which visual model applies (e.g. 1, 2 or 3). For analyses with different animal visual systems, we recommend separate analyses rather than making use of this feature.

Note: The different visual models need to have the same spectral sensitivities, channel names and weber fractions. This functionality is aimed to allow to account for differences in lighting. For the modelling of different visual systems, we recommend completing separate batch processing cycles.

#### 4.3 Specifying the ROIs

After having opened the image using the 'Generate multispectral image' function and having rotated the image, transform the image to a cone catch image using the 'Convert to cone catch' function. This will allow checking for correct cone mapping. Alternatively, this step can be omitted. Either way, to obtain the ROIs we recommend working with a colour image rather than a grey-scale image. It is often easier to see animals and objects that way. It is important that the ROIs are obtained with the image being rotated into its final position. To obtain an RGB colour image, go to 'Make presentation image' and tick the 'Convert to RGB box'. Use the 'Image' -> 'Adjust' -> 'Brightness & Contrast' -> 'Auto' (or manually if needed) to make the animal as visible as possible. Don't worry about other parts of the image getting too bright. This is just to see the outline of the animal as clearly as possible.

First, set the size standard by selecting the 'line tool' to draw a line along the length of your size standard & press 'S' to give its units and save the ROI. We recommend working in mm, i.e. 1cm = 10.

Select the outline of the 'animal' using the 'Freehand selection' tool. Use the zoom function ('Ctrl' + 'alt' + mouse wheel or zoom buttons) to get a nice and close look at the outline of the animal as above. Use the 'Brush Selection Tool' (double click to set size) with 'alt' to add to ROI and 'Shift' to subtract from the ROI to refine the ROI. Press 't' to save the ROI once happy and rename it as 'Animal' (case sensitive). Make sure the 'Show all' box in the ROI manager is disabled.

Select the 'animal+background' ROI by roughly outlining the area of interest <u>including</u> the animal. Press 't' and then 'rename' to 'animal+background' (note: case sensitive!). Avoid areas with excessive shadows cast by the lighting or colour standard. This step is important to exclude unwanted parts of the image.

Combine 'animal+background' with 'animal' to create the final 'background' ROI that does not contain the animal. Use 'CTRL' to select both ROIs in the ROI manager.

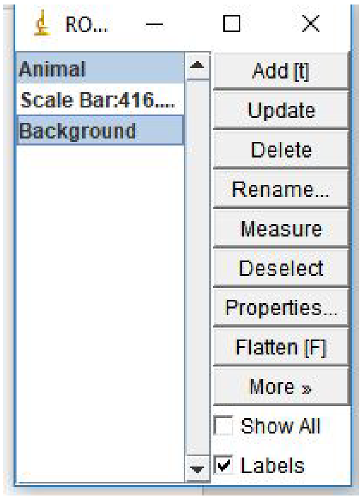

Go to 'More' -> 'XOR'. This will subtract the animal from the background. Press 't'. Rename the new ROI to 'background'. Check all ROIs by selecting them in the ROI manager and observing the ROI in the image. If there are areas that need to be excluded from the background use the same technique as for the 'animal' to remove these areas and save the modified ROI. Save the ROIs once you are happy.

#### 5. Using the batch script

To launch the QCPA batch script, go to 'Plugins' -> 'micaToolbox' -> 'Batch QCPA' and select the directory containing your prepared dataset. For general considerations on parameter choices, please see the electronic supplement of the original QCPA paper, as well as the website.

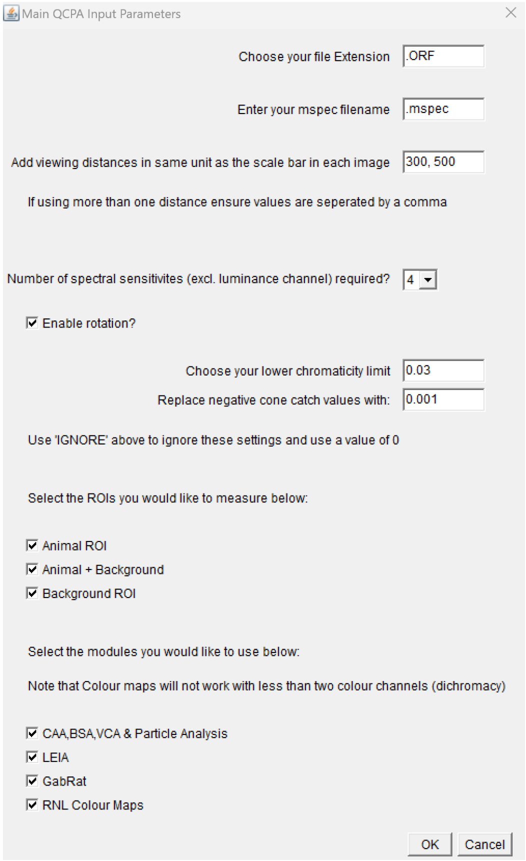

1. Specify the format of your RAW images
2. If you have changed the name of your .mspec files, specify the name here. This name needs to be identical in all images. We recommend not saving a specific name when creating the files.
3. Choose the desired viewing distance(s). The closer the distance and higher the visual acuity, the longer image processing will take.
4. Choose the number of colour channels in your visual system.
5. Select if the images should be rotated or not. We recommend doing so.
6. Choose your lower chromaticity limit.
7. Choose the replacement value for negative cone catch estimates.
8. Select the ROIs you would like to measure. Note: The script assumes all three ROIs to be present. If you want to apply the script to datasets without animals or objects you might want to try creating empty or tiny ROIs.
9. Select the modules you would like to run.

#### 5.1 'CAA, BSA, VCA & Particle Analysis' settings

1. Specify Weber fractions. These will be used for the clustering and the RNL ranked filter. Make sure that these names match the channels specified in your visual model (no need to match the order though!). The luminance channel is automatically detected as the last row in your visual model and will be called 'lum' irrespective of the name given in your model (e.g. dbl or mw).

   Note: Your visual models will need to have a luminance channel. To add one, add it to the spectral sensitivity file, save the file and redo your visual model.

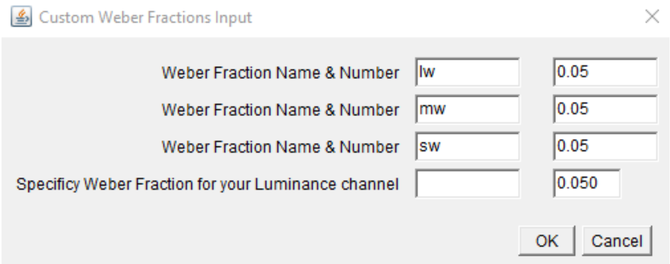

2. Specify your desired Acuity settings. Unless your ROIs are rectangular, use 'Gaussian'.
3. Select if you want to use the RNL ranked filter or not. We recommend the use of RNL ranked filtering to compensate for artefacts from the acuity modelling.

4. Choose your acuity modelling settings. We recommend using the defaults displayed here in combination with your required cycles per degree acuity value.

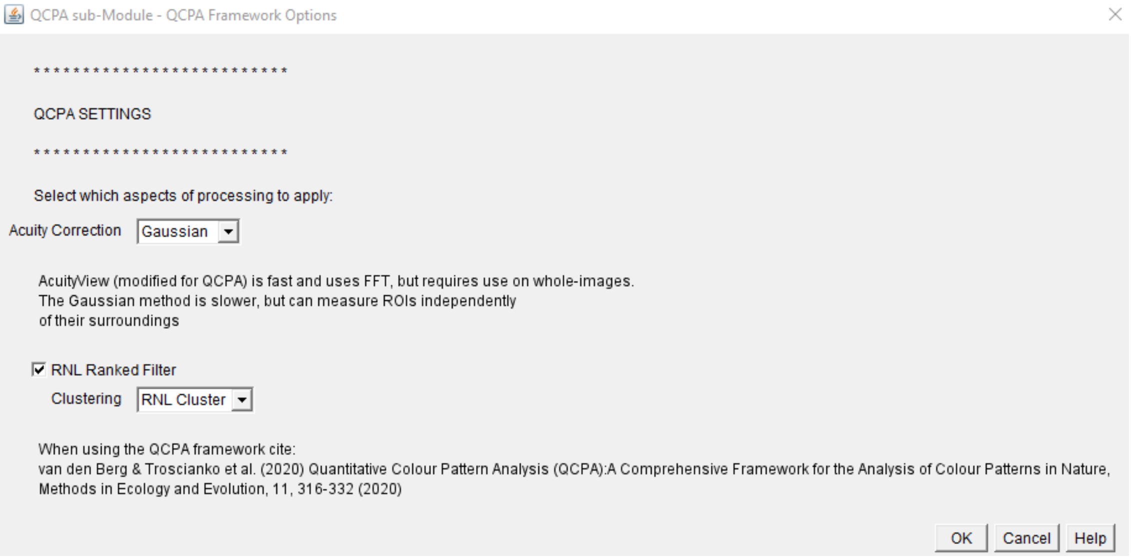

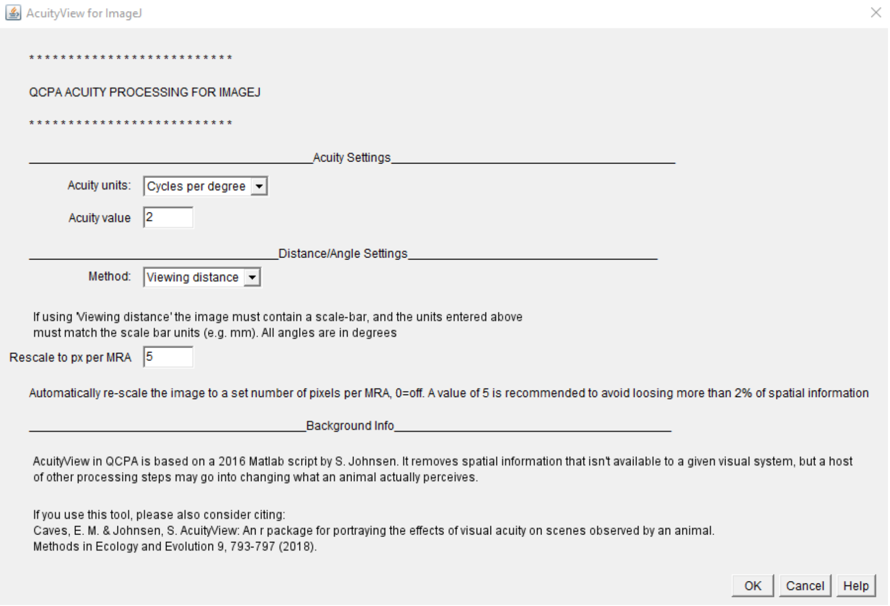

5. Choose your RNL ranked filter options.

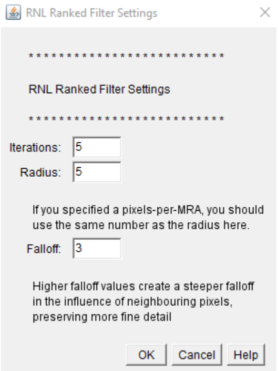

6. Choose your RNL clustering settings. Note that Naïve Bayes clustering requires manual input and is not available in the batch script.

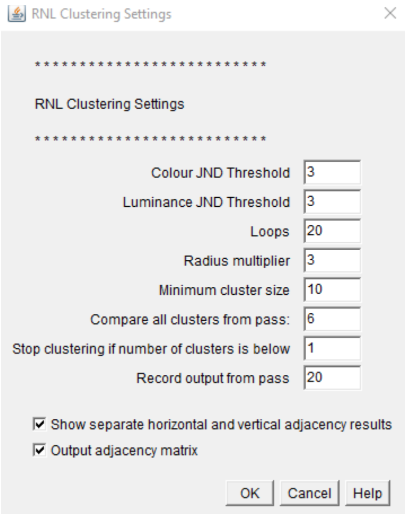

7. Select the appropriate visual models. Note that these need to match ordering specified in your .txt files. That is, model #1 needs to be '1' in your .txt files. The interface shows both, the .java and the .class files for each model. Either one works fine.

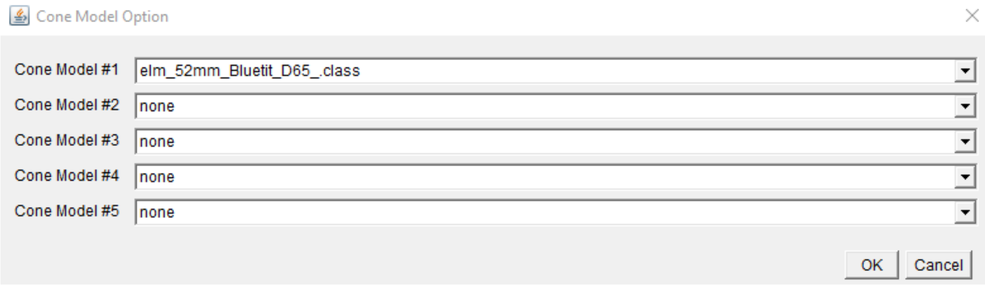

8. Sit back and enjoy! All outputs and log files will be saved in your data next to your images. Please provide the log files in the electronic supplement or data of any publications for reproducibility.

#### 5.2 'LEIA' settings

1. If the user selects only LEIA to be applied, then the following additional input is required. Fore using LEIA, we recommend using the RNL ranked filter but not clustering. If LEIA is selected in addition to other QCPA modules, the user will provide a unique set of parameter choices for either.

   The user has the choice to log-transform the LEIA output. This might provide additional insights as many behaviours might respond to the log intensity of visual input. The user has the option to visually analyse horizontal and vertical LEIA edge contrast in the image separately and set the upper intensity level of the LEIA images stored in the LEIA folder of each image.

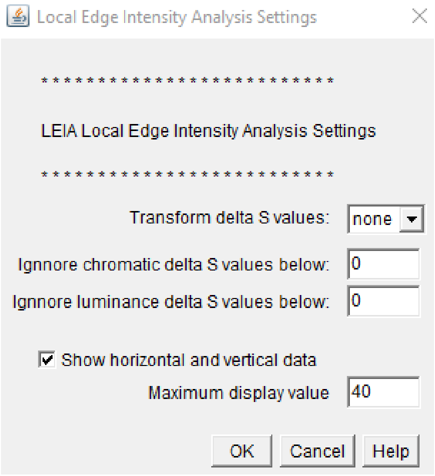

2. Sit back and enjoy! All outputs and log files will be saved in your data next to your images.

#### 5.3 'GabRat' settings

1. The user interface will ask for an acuity value if GabRat is selected by itself or in combination with other modules requiring acuity modelling. However, unlike other modules considering the 'maximal' acuity, GabRat is recommended to be run at the resolution (and thus spatial frequencies) where a visual system is best at resolving spatial detail. Therefore, the acuity value provided in this window will override the previously supplied acuity value.

   Gabrat will by default use the 'Animal' ROI to define the outline of the Animal. Therefore GabRat output for the 'Background' and 'Animal + Background' ROIs are of no scientific value as GabRat measures along the outline of an animal or object. We recommend GabRat to be run without RNL ranked filtering or clustering. If GabRat is run in combination with other modules, each will request its own set of parameters.

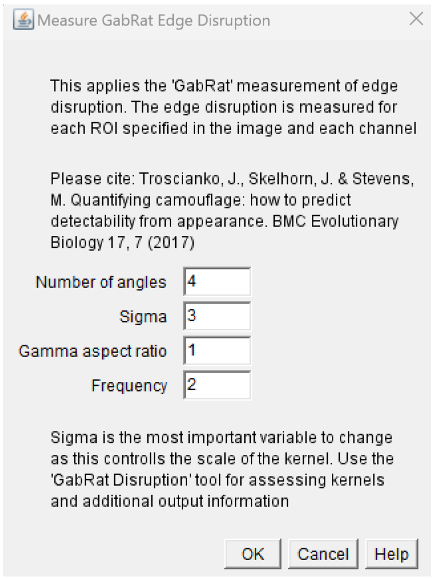

2. Sit back and enjoy! The script will save a .csv file containing the GabRat values for each channel and ROI.

#### 5.4 'Colour Maps' settings

1. When running the Colour map module, the script will ask you to specify the resolution of the colour maps in addition to any input needed for RNL filtering or clustering. We recommend applying RNL filtering but not clustering when using this tool. If the Colour Maps module is used in combination with other QCPA modules, each will request its own set of inputs.

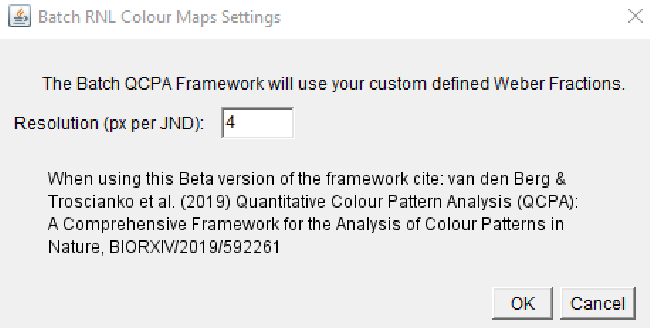

2. Sit back and enjoy! The script will save the colour maps and overlay analysis for each ROI with your images. If you want to display the colour maps, this will have to be done manually. The script will not provide you with colour maps for each particle in a ROI. If you want to get colour maps for individual particles (read: colour pattern elements), we recommend clustering your image and run colour maps on the RNL filtered image using the ROI list obtained from clustering. This will have to be done manually.
