## Supplementary material for "Automated Workflows using Quantitative Colour Pattern Analysis (QCPA): A Guide to Batch Processing and Downstream Data Analysis": Worked examples (from another version of this preprint: version 2, 2023-11-10)

### Example 1: A Trichromatic observer without UV sensitivity

This example uses data of nudibranch molluscs photographed underwater. Nudibranchs are incredibly diverse, colourful marine animals known for their defensive colouration. This is actual data from ongoing research.

We will use the visual system of a triggerfish (*Rhinecanthus aculeatus*) to analyse the images with all available analyses. This example highlights the use of two different visual models, as we pretend some images in the test data were taken with a different light source. We will base our modelling on the cone responses under a slightly green underwater illumination at 5m depth.

To run this example, you will need:

1. The test data provided here: https://doi.org/10.48610/3cdcc1f
2. The following visual models (provided with the test data):
    a. Olympus_PEN_E_PL5_Olympus_60mm_f2_8_PV62white_VK6R_Combined_to_Triggerfish_5m_green_water
    b. Olympus_PEN_E_PL5_Olympus_60mm_f2_8_VK6R_to_Triggerfish_5mGreenWater
3. The latest version of the MICA toolbox & ImageJ available here
4. The correctly installed QCPA batch script, available here: https://github.com/cedricvandenberg/QCPA-batch-script

Step 1: Launch the batch script (Plugins -> micaToolbox -> Batch QCPA -> Launch Batch QCPA)

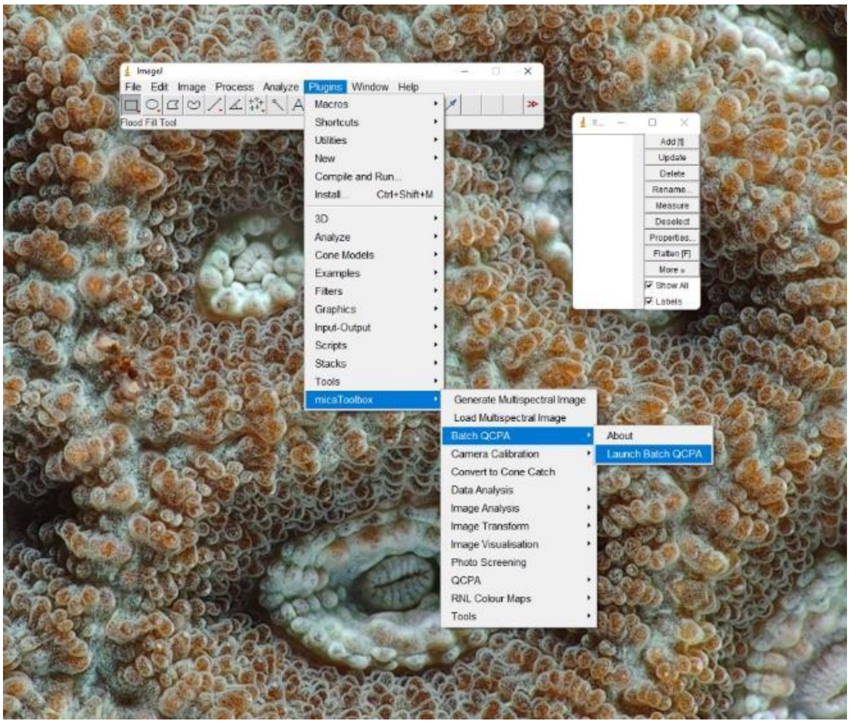

Step 2: Select the 'Test Data' folder in the 'Tri-chromat no UV' example folder.

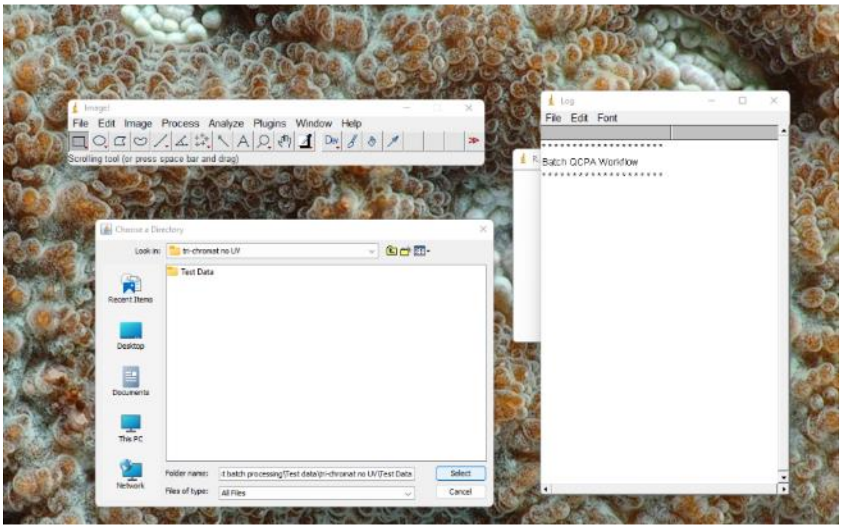

Step 3: Provide the general input setting for the analysis.

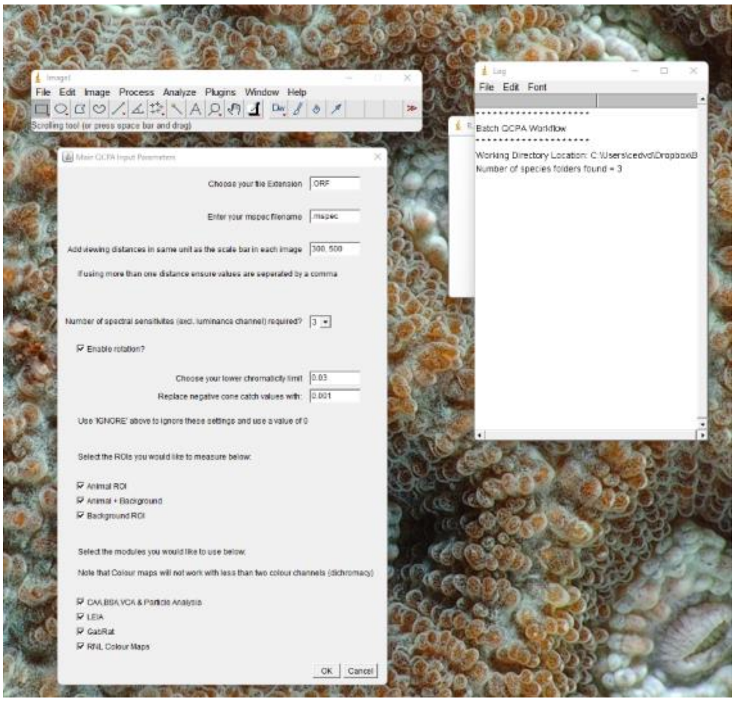

a. We will be working with '.ORF' images, the RAW file format for Olympus cameras.
b. We have not renamed our .mspec images. Thus, the script will look for files simply called '.mspec'.
c. We will simulate viewing distances of 30 cm and 50 cm. This will make processing comparably fast. Note that, the larger the images, the higher the spatial acuity of the modelled visual system and the closer the viewing distance, the longer processing will take as the resulting images will remain large.
d. Triggerfish (*Rhinecanthus aculeatus*) have 3 known cone types and thus, spectral sensitivities, involved in bright-light colour vision.
e. As the images were taken with variable animal rotations, each animal folder contains a .txt file called 'rotation.txt' specifying the number of degrees the image needs to be rotated to be aligned with the anterior-posterior axis of the animal. Therefore, 'Enable rotation' should be ticked.
f. We will choose a lower chromaticity limit of 0.03. This means that, below 3% luminance, the observer should not be able to perceive any colour.
g. We will replace negative cone catch values in our images with values of 0.001.
h. We will analyse all three ROIs available in the dataset: The animal, the animal and its background and the background by itself.
i. We will apply all modules available in the QCPA batch script.

Step 4: Provide Weber fractions

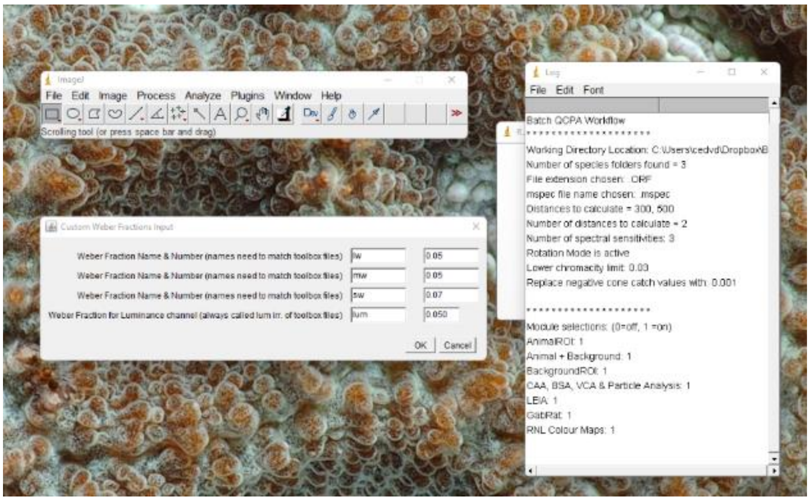

*a.* Studies using *R. aculeatus* as a modelled observer typically assume a very conservative noise level of 0.05 in every type of photoreceptor. Luckily, we have a good understanding of the relative abundance of each type of photoreceptor and we can calculate the corresponding channel-specific Weber fractions: lw: 0.05, mw: 0.05, sw: 0.07.

*b.* *R. aculeatus* uses its double cone (the fused 'unit' of the mw & lw receptors) for luminance contrast detection. Thus, in the csv file in the toolbox that was used to train the mapping function, the luminance channel is called 'dbl'. Therefore, the luminance channel in the mapping function is also called 'dbl'. However, the batch script automatically assumes the last channel in that file to be the luminance channel and will refer to it as 'lum'. Thus, no need to call it anything else and, importantly, make sure your sensitivity file in the toolbox, used to train your mapping function, has a luminance channel.

Step 5: Provide settings relating to module 1 – CAA, BSA, VCA and Particle Analysis

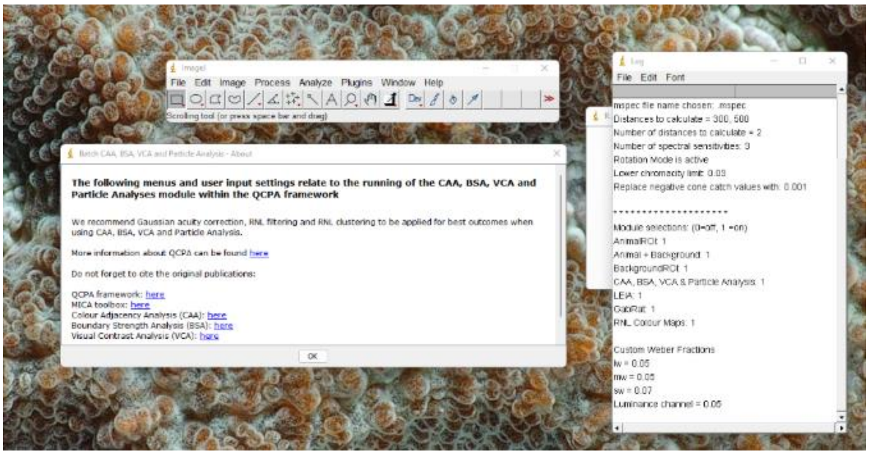

a. The script will come up with an intro panel with useful links and recommendations for settings and reminders to cite the correct corresponding literature for use in publications.

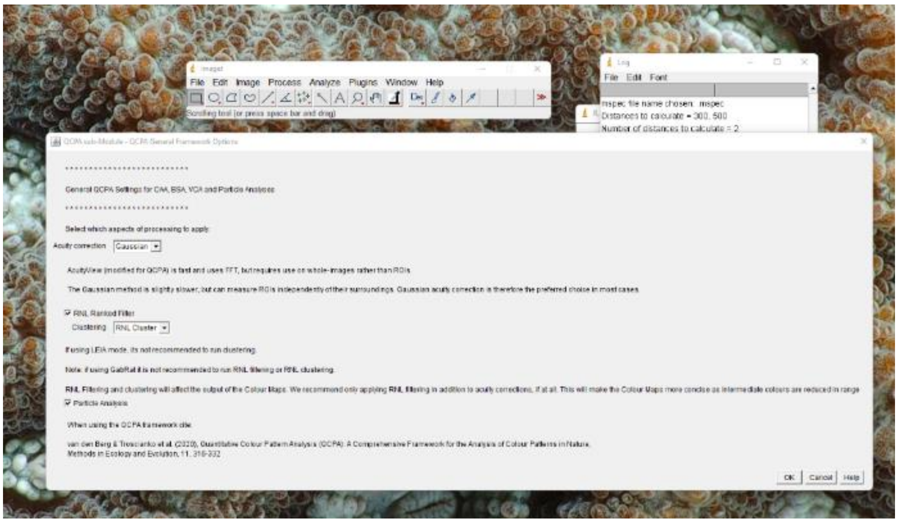

b. We will be using the Gaussian acuity modelling (not AcuityView) implemented in the QCPA as the outlines of our animals and the backgrounds are not rectangular and we are interested in analysing ROIs individually, rather than the whole image.
c. To use module 1, clustering is mandatory. We, therefore, use the RNL clustering, as the naïve Bayes clustering is not available because it would require manual input for each image.
d. We will enable particle analysis
e. RNL filtering is enabled, as recommended following spatial acuity modelling.

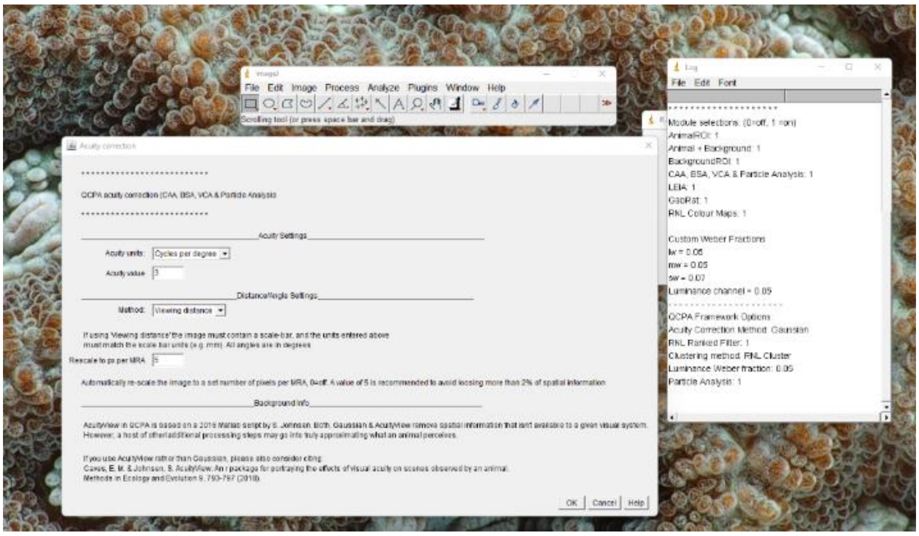

f. We will use acuity provided in cycles per degree (cpd). Behavioural experiments and histological data suggest a maximum acuity of about 3-5 cpd in triggerfish. We'll be conservative and use 3 cpd.
g. We will be using the 'viewing distance' method, which is the only option available in the batch script.
h. We will rescale our images to a pixel/MRA ratio of 5 to maximise processing speed while minimising the loss of relevant spatial information.

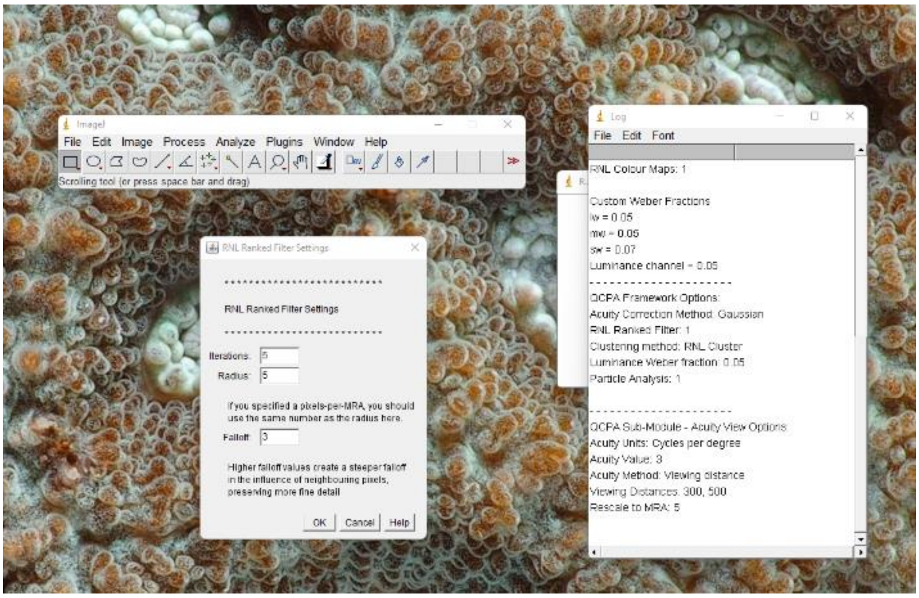

i. We will be using the RNL filter with 5 iterations, a radius of 5 and a falloff of 3.

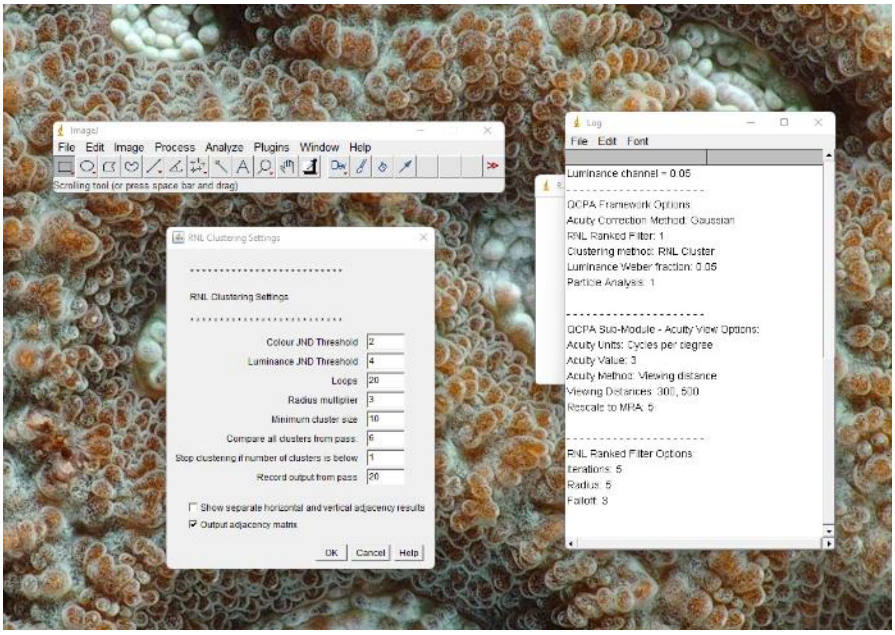

j. We will be running the analysis with a chromatic JND threshold of 2 ΔS and an achromatic threshold of 4 ΔS. These values align with recent findings from behavioural experiments.
k. The rest of the settings we will leave at default. However, we will tick the options for all the output.

<u>Step 6:</u> Provide input settings relating to module 2 - LEIA

Most settings for LEIA will be similar to module 1. However, for LEIA, we will:

a. NOT cluster the images, as we want to keep as many informative edges as possible.
b. NOT run particle analysis, as our images for LEIA will not be clustered.

All other settings are identical.

LEIA requires a set of unique input choices at the end:

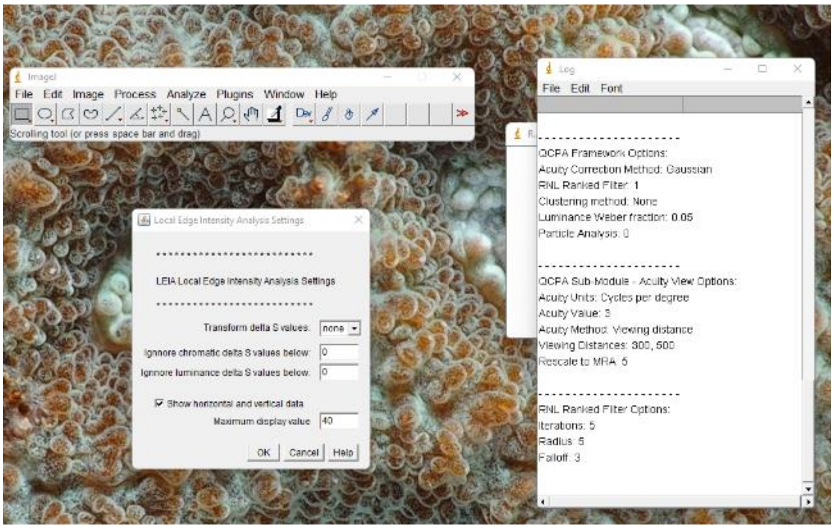

a. Well not transform the ΔS values of the edges detected by LEIA.
b. We will ignore chromatic and achromatic edges with negative values
c. We will choose to have the horizontal and vertical edges reported in addition to the overall edge contrast.
d. We will choose a maximum display contrast of 40 (this has no impact on the data).

<u>Step 7:</u> Provide input settings relating to module 3 - GabRat

Most settings for GabRat will be similar to the other modules. However, for GabRat we will:

a. NOT cluster the images
b. NOT run particle analysis, as our images for GabRat will not be clustered
c. NOT run RNL clustering, as we want edges in our images left as unprocessed as possible following acuity modelling.
d. Run our Gaussian acuity correction with a value of 1cpd, the assumed λmax of the CSF in a triggerfish (this is unique to GabRat).

GabRat requires a set of specific inputs at the end:

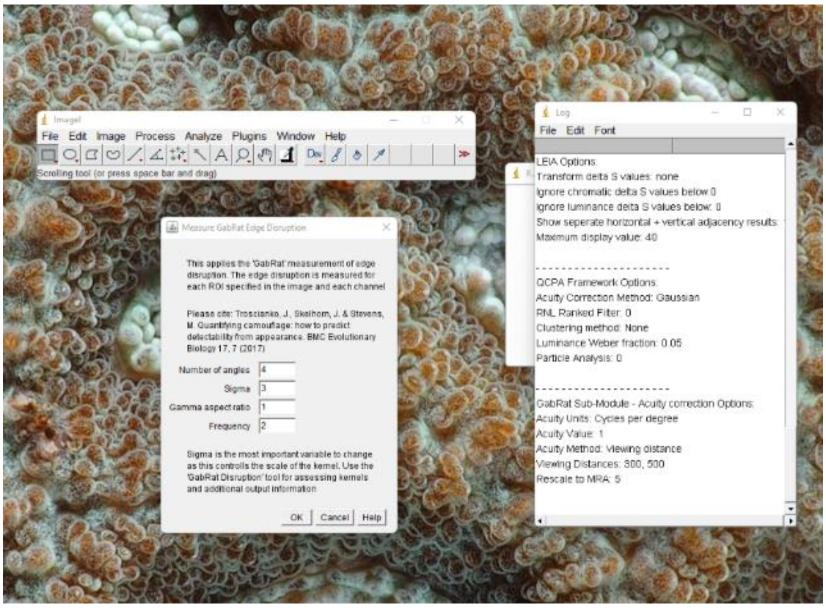

We will leave these settings at default. In a nutshell: The acuity modelling applied to the image means we won't need to adjust the properties of the Gabor filter (leave default settings) → It's all happening at the scale of a receptive field already, due to our image rescaling.

<u>Step 8:</u> Provide input settings relating to module 4 – Colour Maps

Most settings for Colour Maps will be similar to the other modules. However, for Colour maps we will:

a. NOT cluster our images
b. Use the RNL filter, as we want to restrict the range of colours in our image (try with and without to see the difference).
c. Not enable particle analysis, as our images won't be clustered.

Colour Maps require the user to set the resolution of the Colour Map files. We recommend using the default resolution of 4 pixels per JND.

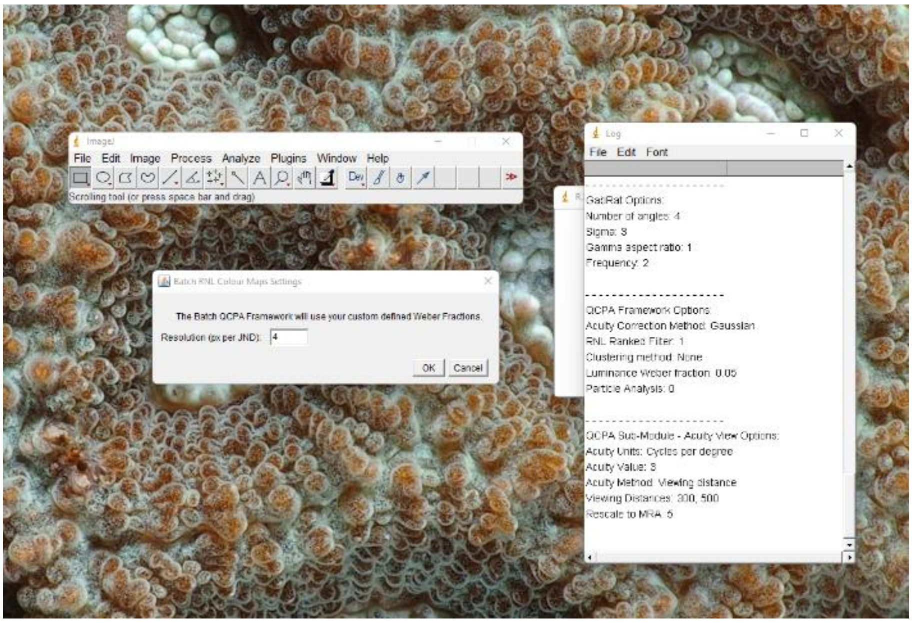

<u>Step 9:</u> Define which cone mapping functions ('visual models') to use

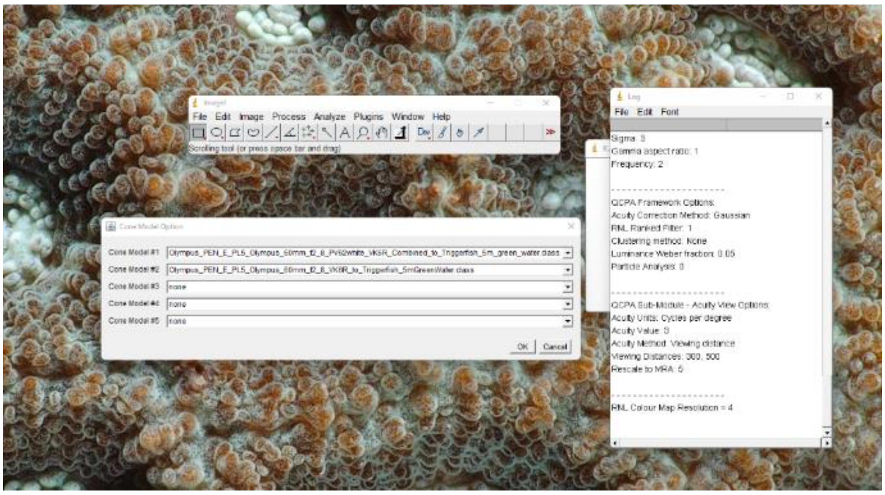

a. We will be using: Olympus_PEN_E_PL5_Olympus_60mm_f2_8_PV62white_VK6R_Combined_to_Triggerfish_5m_green_water as our cone model number 1, and Olympus_PEN_E_PL5_Olympus_60mm_f2_8_VK6R_to_Triggerfish_5mGreenWater as our model number 2.
b. You might notice that, in fact, all images in the dataset have model 1 specified in their corresponding .txt file, so the analysis would run even if we had just specified one model. We're doing this here just to demonstrate how.

The analysis should take a couple of minutes to run.

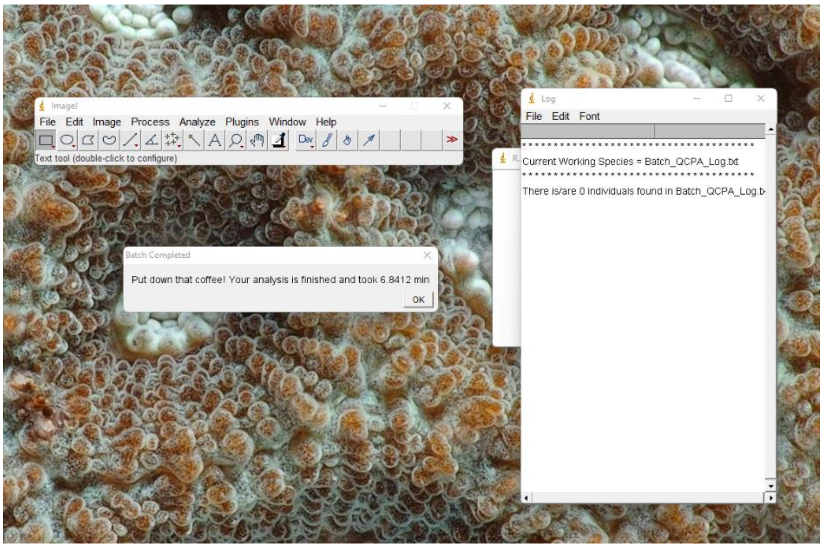

### Example 1: Data Output

The batch script will provide a detailed log for the settings used at the top level. Please upload this log with any published data and modelling files to promote reproducibility.

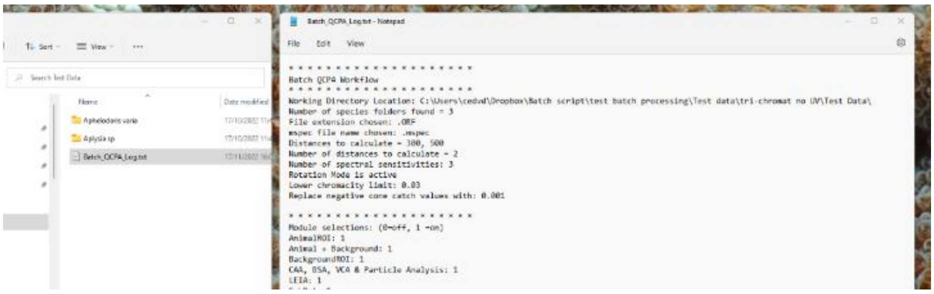

All data output will be stored in a distance-specific folder. In our case 30cm and 50cm.

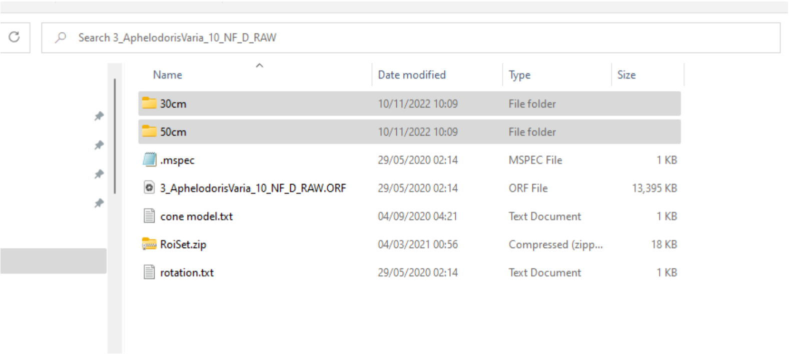

Within each distance folder, a log file for each ROI and module can be found, together with all generated output **for module 1**:

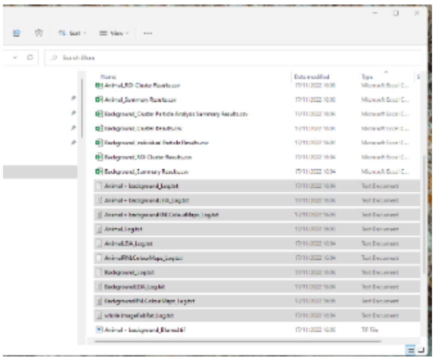

Specifically, for each ROI you will find:

'ROI NAME'_blurred.tiff → a multispectral .tiff of your ROI after acuity modelling & RNL filtering (or without filtering if you don't use the RNL filter). This is great for visualisations (i.e. false colour presentation images).

'ROI NAME'_Cluster Particle Analysis Summary Results.csv → A summary of all individual clusters (summarised across individual sub particles) in the segmented image. This is a great way to get a suite of descriptors summarising each colour pattern element, such as its average orientation, total size, solidity, etc. This is a base ImageJ function documented here.

'ROI NAME' Cluster Results.csv → A range of basic summary metrics for each cluster in the ROI, such as different mean chromaticity metrics, mean receptor channel stimulation etc.

'ROI NAME'_Clustered.tiff → A multispectral .tiff of your ROI after RNL clustering. This is great for visualisations (i.e. false colour presentation images).

'ROI NAME'_Individual Particle Results.csv → Particle descriptors for all sub-particles. A good source of information on shape of individual colour pattern elements.

'ROI' Log.txt → A log file of all processing steps and their respective settings applied in module 1.

'ROI NAME'_ROI Cluster results.csv → Summary of the spectral properties of the colour pattern elements. This is where you find the transition matrices. This file is the basis for all computations for the CAA, BSA, VCA.

‘ROI Name’_Summary Results.csv → All CAA, VCA, BSA parameters.

‘ROI NAME’_Zonemap.tiff → A .tiff file of the clustered ROI as a zone map, i.e. with each cluster numbered 1 to k.

**For modules 2 – 4**, all data output is stored in corresponding subfolders:

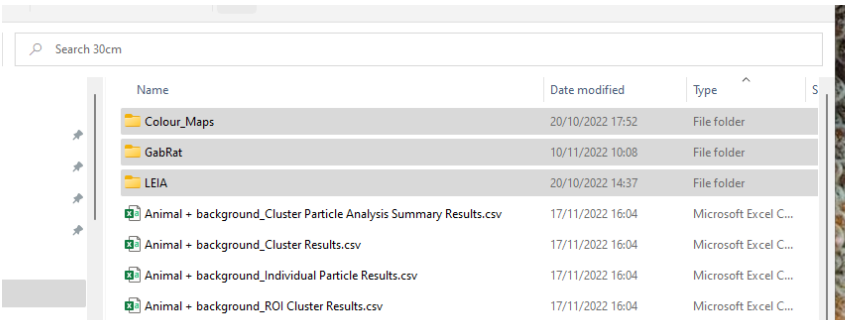

**Colour Maps:**

Each ROI is saved as a colour map cloud .tiff file that can be used to visualise colour space and/or calculate colour space overlap using the colour map functions in QCPA (see here).

Additionally, a detailed log can be found in the main folder.

**LEIA:**

For each ROI, the user will have the numerical output of LEIA in a .csv file as well as the visualised chromatic and achromatic LEIA contrast saved as a .tiff. The latter are fantastic for visualisation purposes.

Additionally, a detailed log can be found in the main folder.

**GabRat:**

Here, the user will find a .csv file with the GabRat values for all ROIs. Note that only the animal ROI is of significance, as the kernel runs along the outside of the animal ROI.

Additionally, a detailed log can be found in the main folder.

To compile data out of these folders & files, please see the R-script library provided here:

https://github.com/CaraConradsen/QCPA-r-script.

### Example 2: A Tetrachromatic observer with UV sensitivity

For this example, we will investigate a bird predator (European Bluetit, *Parus major*) looking at spiders (*Tamopsis brisbanensis*) on a tree trunk. These image data and calibration files have been kindly provided by Alfonso Aceves.

To run this example, you will need:

1. The test data provided here: https://doi.org/10.48610/3cdcc1f
2. The following visual model (provided with the test data):
    a. Nikon_D7000_Novoflex_35mm_D65_to_Bluetit_D65
3. The latest version of the MICA toolbox & ImageJ available here
4. The correctly installed QCPA batch script, available here: https://github.com/cedricvandenberg/QCPA-batch-script

Step 1: Launch the batch script (Plugins -> micaToolbox -> Batch QCPA -> Launch Batch QCPA)

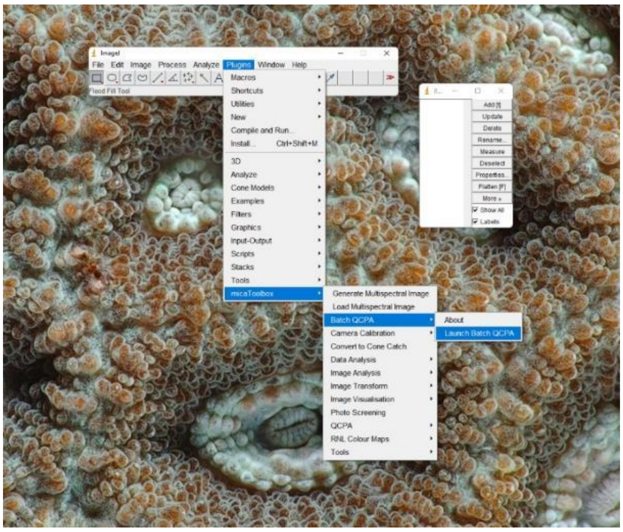

Step 2: Select the 'Test Data' folder in the 'tetra-chromat with UV' example folder.

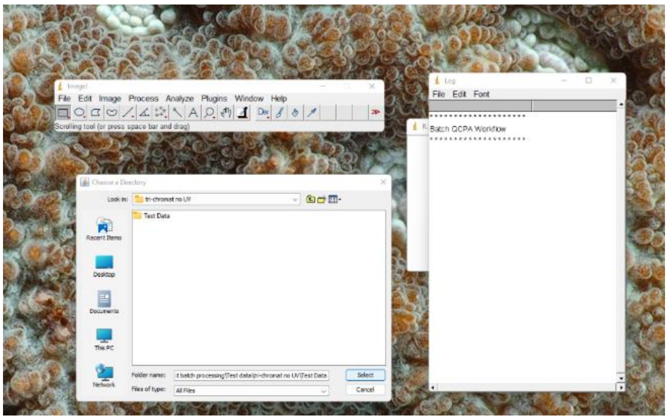

<u>Step 3:</u> Provide the general input setting for the analysis.

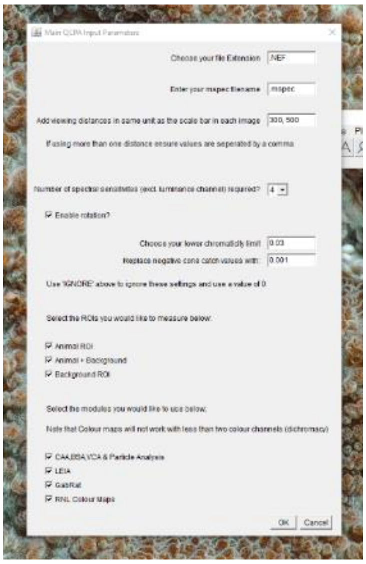

a. We will work with '.NEF' images, the RAW file format for Nikon cameras.
b. We have not renamed our .mspec images. Thus, the script will look for files simply called '.mspec'.
c. We will simulate viewing distances of 30 cm and 50 cm. This will make processing comparably fast. Note that, the larger the images, the higher the spatial acuity of the modelled visual system and the closer the viewing distance, the longer processing will take as the resulting images will remain large.
d. Bluetits (*Parus major*) have 4 known cone types and thus, spectral sensitivities, involved in bright-light colour vision, plus a double cone that is responsible for luminance contrast perception.
e. As the images were taken with variable animal rotations, each animal folder contains a .txt file called 'rotation.txt' specifying the number of degrees the image needs to be rotated to be aligned with the anterior-posterior axis of the animal. Therefore, 'Enable rotation' should be ticked.
f. We will choose a lower chromaticity limit of 0.03. This means that, below 3% luminance, the observer should not be able to perceive any colour.
g. We will replace negative cone catch values in our images with values of 0.001.
h. We will analyse all three ROIs available in the dataset: The animal, the animal and its background and the background by itself.
i. We will apply all modules available in the QCPA batch script.

Step 4: Provide Weber fractions

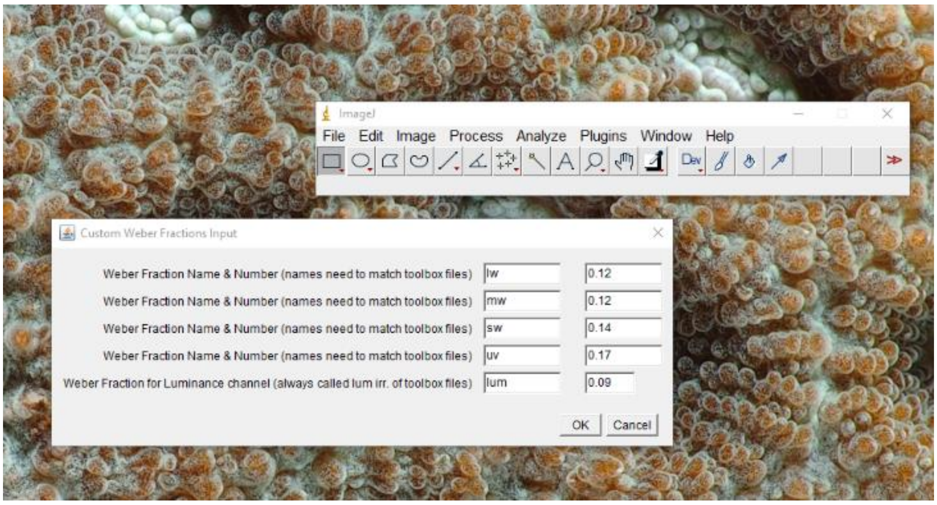

a. Hart et al., 2000 showed that photoreceptors are distributed at the following ratios (uv:sw:mw:lw:dbl): 1:1.92:2.68:2.7 with the (double) dbl cone relative abundance of 4.86. When normalised to the most abundant class of receptors, we get: 0.21:0.4:0.55:0.55:1. Silvasti et al., (2021) have determined that a Weber fraction of 0.2 reflects a 1 ΔS JND for the lw cone and thus, following the equations outlined here, we get a receptor-specific noise of 0.09, this then let's us calculate the appropriate Weber fractions as follows: 0.17:0.14:0.12:0.12:0.09 (uv:w:mw:lw:dbl). Here, dbl, is the luminance channel.
b. *P. major* uses its double cone for luminance contrast detection. Thus, in the csv file in the toolbox that was used to train the mapping function, the luminance channel is called 'dbl'. Therefore, the luminance channel in the mapping function is also called 'dbl'. However, the batch script automatically assumes the last channel in that file to be the luminance channel and will refer to it as 'lum'. Thus, no need to call it anything else and, importantly, make sure your sensitivity file in the toolbox, used to train your mapping function, has a luminance channel.

Step 5: Provide settings relating to module 1 – CAA, BSA, VCA and Particle Analysis

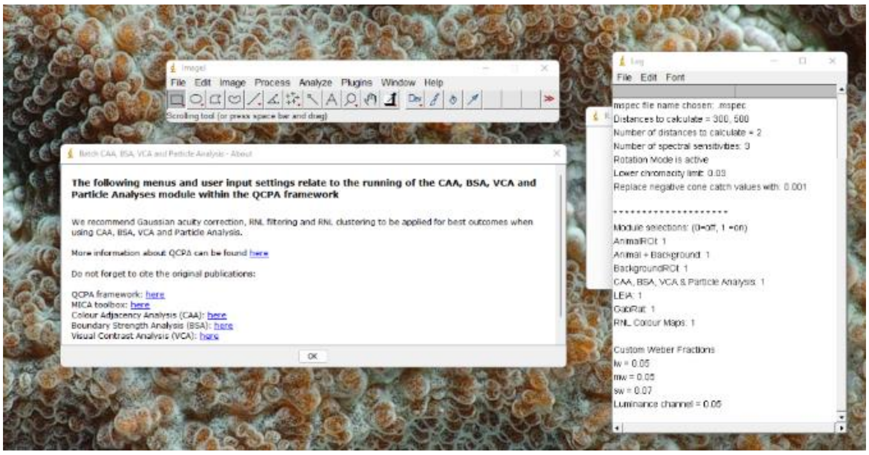

a. The script will come up with an intro panel with useful links and recommendations for settings and reminders to cite the correct corresponding literature for use in publications.

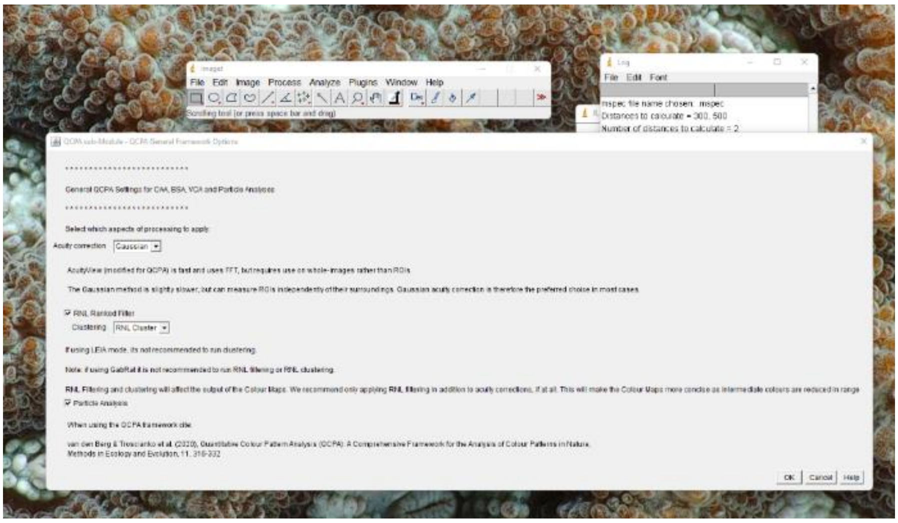

b. We will be using the Gaussian acuity modelling (not AcuityView) implemented in the QCPA as the outlines of our animals and the backgrounds are not rectangular and we are interested in analysing ROIs individually, rather than the whole image.
c. To use module 1, clustering is mandatory. We, therefore, use the RNL clustering, as the naïve Bayes clustering is not available because it would require manual input for each image.
d. We will enable particle analysis
e. RNL filtering is enabled, as recommended following spatial acuity modelling.

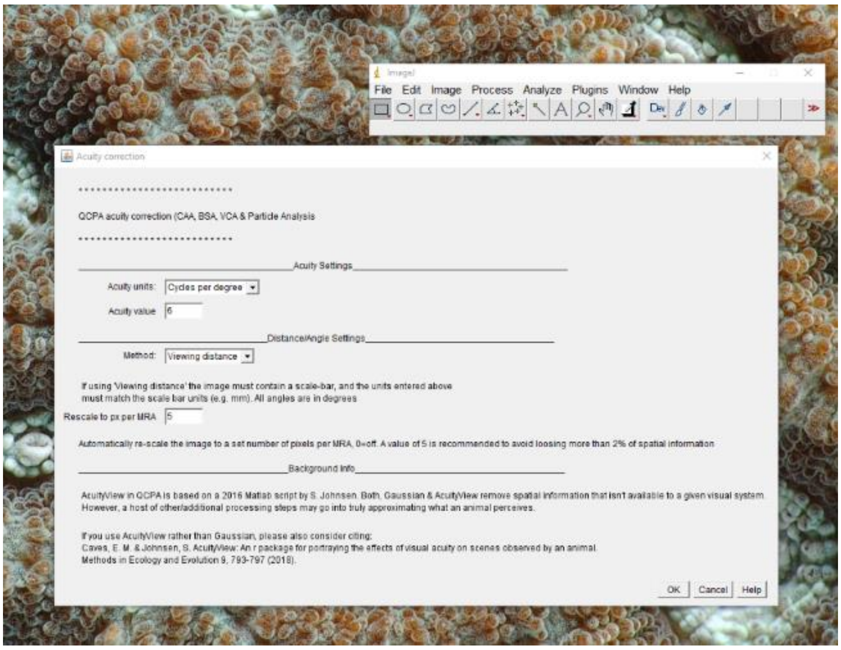

f. We will use acuity provided in cycles per degree (cpd). Behavioural experiments and histological data suggest a maximum acuity of about 6 cpd in tits (Martin, 2018) which we will use here.
g. We will be using the 'viewing distance' method, which is the only option available in the batch script.
h. We will rescale our images to a pixel/MRA ratio of 5 to maximise processing speed while minimising the loss of relevant spatial information.

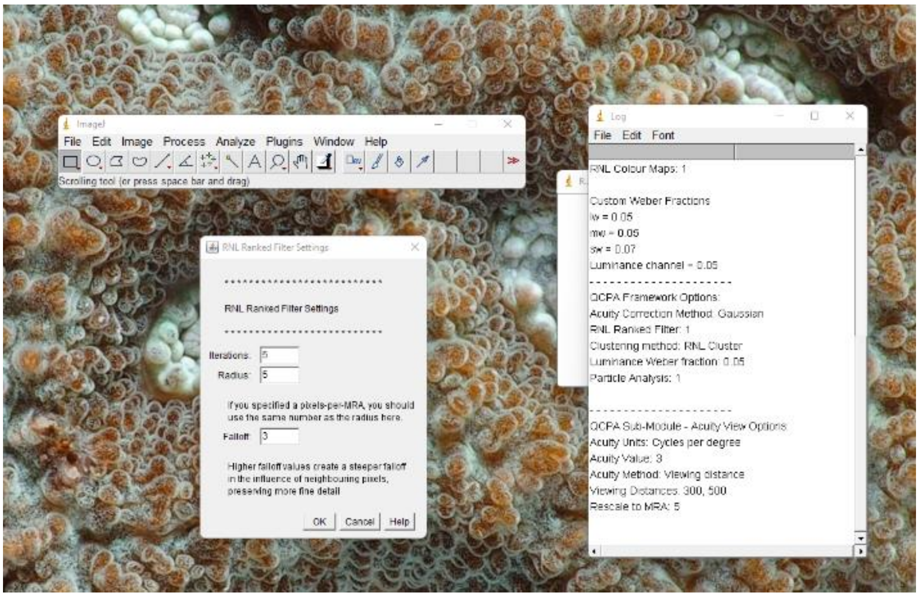

i. We will be using the RNL filter with 5 iterations, a radius of 5 and a falloff of 3.

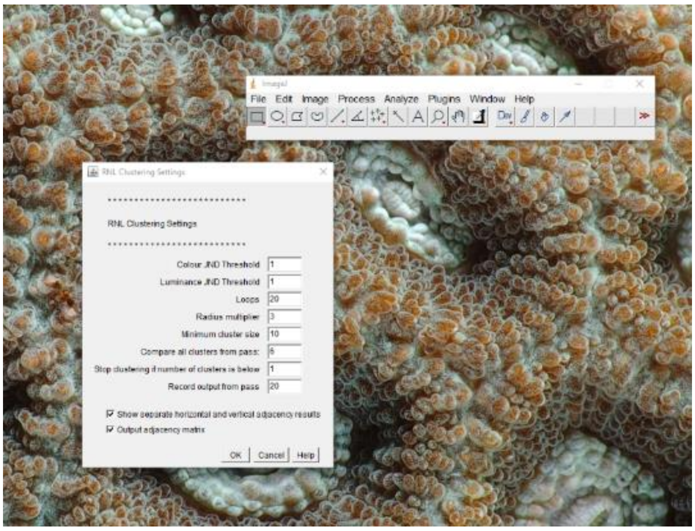

j. We will be running the analysis with a chromatic JND threshold of 1 ΔS and an achromatic threshold of 1 ΔS. These values are much less conservative as we have calibrated our weber fractions using findings from behavioural experiments.
k. The rest of the settings we will leave at default. However, we will tick the options for all the output.

<u>Step 6:</u> Provide input settings relating to module 2 - LEIA

Most settings for LEIA will be similar to module 1. However, for LEIA, we will:

a. NOT cluster the images, as we want to keep as many informative edges as possible.
b. NOT run particle analysis, as our images for LEIA will not be clustered.

All other settings are identical.

LEIA requires a set of unique input choices at the end:

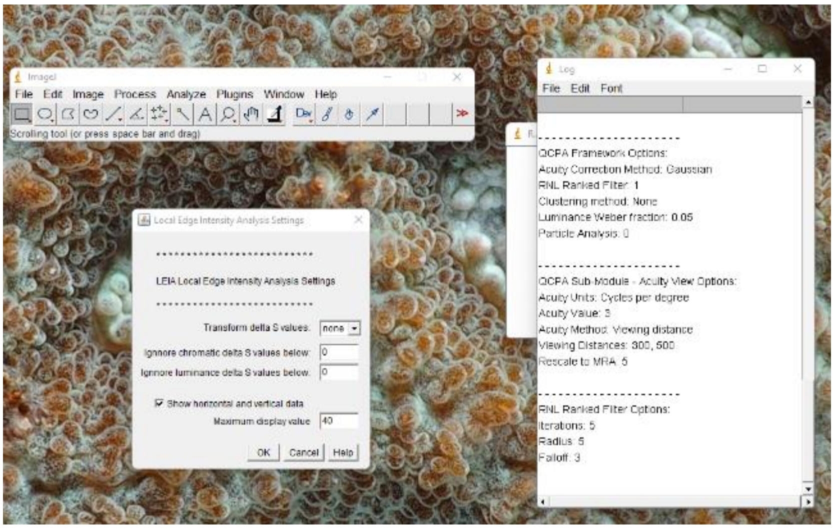

a. Well not transform the ΔS values of the edges detected by LEIA.
b. We will ignore chromatic and achromatic edges with negative values
c. We will choose to have the horizontal and vertical edges reported in addition to the overall edge contrast.
d. We will choose a maximum display contrast of 40 (this has no impact on the data).

<u>Step 7:</u> Provide input settings relating to module 3 - GabRat

Most settings for GabRat will be similar to the other modules. However, for GabRat we will:

a. NOT cluster the images
b. NOT run particle analysis, as our images for GabRat will not be clustered
c. NOT run RNL clustering, as we want edges in our images left as unprocessed as possible following acuity modelling.
d. Run our Gaussian acuity correction with a value of 1cpd, the assumed λmax of the CSF in a songbird (Harmening & Wagner, 2011).

GabRat requires a set of specific inputs at the end:

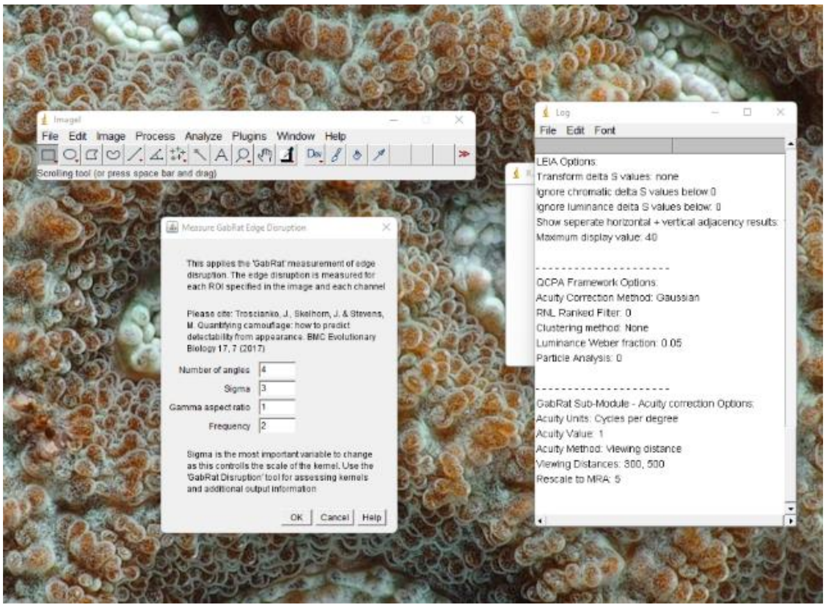

We will leave these settings at default. In a nutshell: The acuity modelling applied to the image means we won't need to adjust the properties of the Gabor filter (leave default settings) → It's all happening at the scale of a receptive field already, due to our image rescaling.

<u>Step 8:</u> Provide input settings relating to module 4 – Colour Maps

Most settings for Colour Maps will be similar to the other modules. However, for Colour maps we will:

a. NOT cluster our images
b. Use the RNL filter, as we want to restrict the range of colours in our image (try with and without to see the difference).
c. Not enable particle analysis, as our images won't be clustered.

Colour Maps require the user to set the resolution of the Colour Map files. We recommend using the default resolution of 4 pixels per JND.

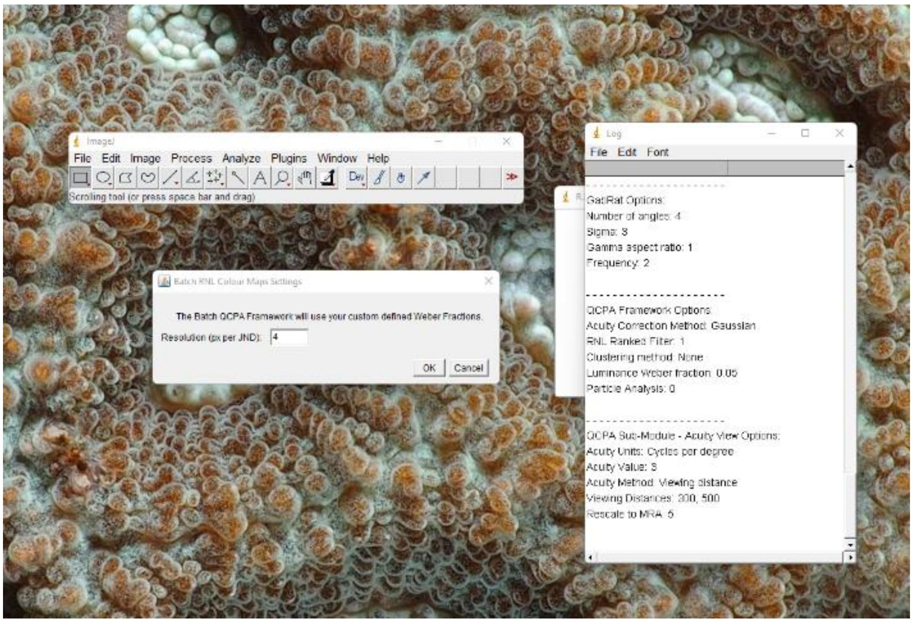

<u>Step 9:</u> Define which cone mapping functions ('visual models') to use

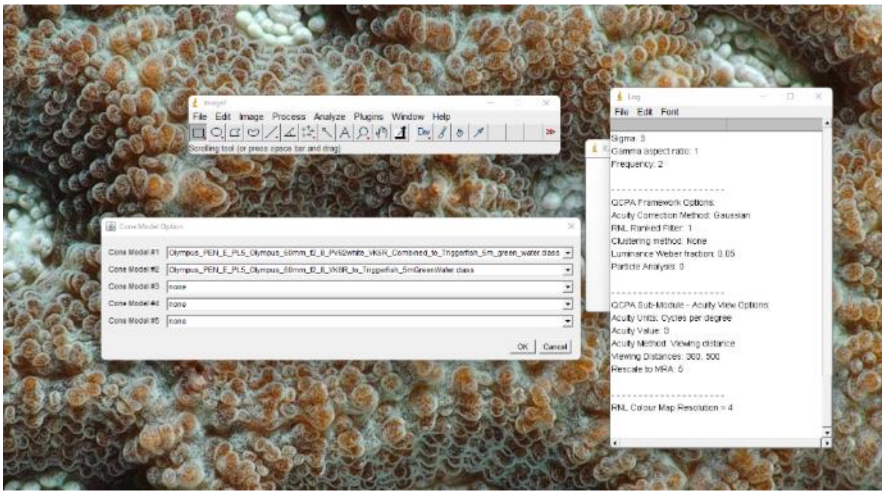

c. We will be using: Nikon_D7000_Novoflex_35mm_D65_to_Bluetit_D65.

The analysis should take a couple of minutes to run. Note that this analysis will take longer than example 1 due to the significantly increased spatial acuity of the viewer.

### Example 2: Data Output

The batch script will provide a detailed log for the settings used at the top level. Please upload this log with any published data and modelling files to promote reproducibility.

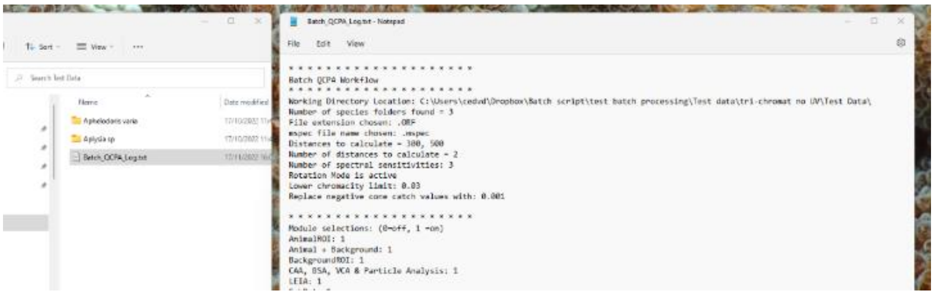

All data output will be stored in a distance-specific folder. In our case 30cm and 50cm.

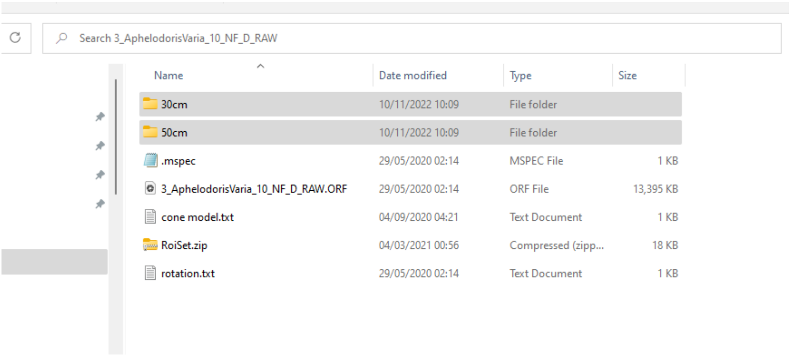

Within each distance folder, a log file for each ROI and module can be found, together with all generated output **for module 1**:

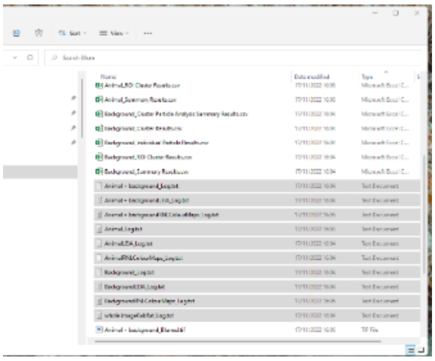

Specifically, for each ROI you will find:

'ROI NAME'_blurred.tiff → a multispectral .tiff of your ROI after acuity modelling & RNL filtering (or without filtering if you don't use the RNL filter). This is great for visualisations (i.e. false colour presentation images).

'ROI NAME'_Cluster Particle Analysis Summary Results.csv → A summary of all individual clusters (summarised across individual sub particles) in the segmented image. This is a great way to get a suite of descriptors summarising each colour pattern element, such as its average orientation, total size, solidity, etc. This is a base ImageJ function documented here.

'ROI NAME' Cluster Results.csv → A range of basic summary metrics for each cluster in the ROI, such as different mean chromaticity metrics, mean receptor channel stimulation etc.

'ROI NAME'_Clustered.tiff → A multispectral .tiff of your ROI after RNL clustering. This is great for visualisations (i.e. false colour presentation images).

'ROI NAME'_Individual Particle Results.csv → Particle descriptors for all sub-particles. A good source of information on shape of individual colour pattern elements.

'ROI' Log.txt → A log file of all processing steps and their respective settings applied in module 1.

'ROI NAME'_ROI Cluster results.csv → Summary of the spectral properties of the colour pattern elements. This is where you find the transition matrices. This file is the basis for all computations for the CAA, BSA, VCA.

'ROI Name'_Summary Results.csv → All CAA, VCA, BSA parameters.

'ROI NAME'_Zonemap.tiff → A .tiff file of the clustered ROI as a zone map, i.e. with each cluster numbered 1 to k.

**For modules 2 – 4**, all data output is stored in corresponding subfolders:

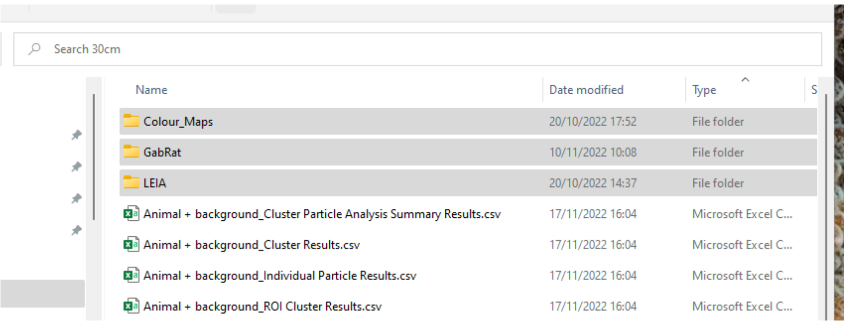

**Colour Maps:**

Each ROI is saved as a colour map cloud .tiff file that can be used to visualise colour space and/or calculate colour space overlap using the colour map functions in QCPA (see here).

Additionally, a detailed log can be found in the main folder.

**LEIA:**

For each ROI, the user will have the numerical output of LEIA in a .csv file as well as the visualised chromatic and achromatic LEIA contrast saved as a .tiff. The latter are fantastic for visualisation purposes.

Additionally, a detailed log can be found in the main folder.

**GabRat:**

Here, the user will find a .csv file with the GabRat values for all ROIs. Note that only the animal ROI is of significance, as the kernel runs along the outside of the animal ROI.

Additionally, a detailed log can be found in the main folder.

To compile data out of these folders & files, please see the R-script library provided here:

https://github.com/CaraConradsen/QCPA-r-script
